# Tau oligomers modulate synapse fate by eliciting progressive bipartite synapse dysregulation and synapse loss

**DOI:** 10.1101/2025.09.19.677215

**Authors:** Kristeen A. Pareja-Navarro, Christina D. King, Grant Kauwe, Yani Y. Ngwala, Doyle Lokitiyakul, Ivy Wong, Aaryan Vira, Jackson H. Chen, Mahima Sharma, Gabriel Navarro, Olfat Malak, Birgit Schilling, Tara E. Tracy

## Abstract

**Background:** Synapse function is critical for cognition, and synapse loss is highly correlated with cognitive decline in Alzheimer’s disease and related dementias. Tau oligomers, which accumulate in the brain in Alzheimer’s disease, can acutely inhibit synaptic plasticity and cause synapse loss. Coordinated presynaptic and postsynaptic function is essential for effective synaptic transmission, and both compartments can be dysregulated by pathogenic tau. However, the series of pathophysiological events triggered by tau oligomers to cause the dysfunction and deterioration of presynaptic terminals and postsynaptic sites remain unclear.

**Methods:** We developed a proximity labeling tool to map the postsynaptic proteome by fusing PSD-95 with APEX2 (APEX2-PSD-95) which was expressed in human induced pluripotent stem cell (iPSC)-derived neurons. We used APEX2-PSD-95 to map the dynamic changes in the postsynaptic proteome with precise temporal resolution after an acute exposure of human iPSC-derived neurons to recombinant tau oligomers for 30 min. Leveraging immunocytochemistry, electrophysiology and electron microscopy, we further delineated the impact of the acute tau oligomer exposure on presynaptic and postsynaptic compartments over time for up to 14 days.

**Results:** The brief exposure of human iPSC-derived neurons to tau oligomers caused a progressive deterioration of synapses, marked by both presynaptic and postsynaptic dysregulation. Postsynaptic proteome mapping revealed an immediate tau oligomer-triggered downregulation of the postsynaptic actin motor proteins Myosin-Va and Myosin-10, which coincided with impaired AMPA receptor (AMPAR) trafficking during synaptic plasticity. This was followed 24 hours later by the upregulation of disease-related proteins, including GSK3β, at postsynaptic sites. The loss of PSD-95-labeled postsynaptic sites at 7 days after tau oligomer exposure preceded the loss of Synapsin-labeled presynaptic terminals at 14 days. The postsynaptic sites that remained exhibited a long-term downregulation of postsynaptic AMPAR levels and sustained synaptic plasticity impairment. Moreover, the remaining presynaptic terminals contained less clusters of vesicles at the presynaptic active zone which was associated with reduced vesicle release probability at synapses.

**Conclusion:** Our findings reveal the series of events underlying tau oligomer-induced bipartite synapse deterioration. The progressive decline of synapses involves the emergence of two synapse fates. One synapse fate involves the persistent weakening of both presynaptic and postsynaptic function, and the other results in synaptic loss.

## Background

Synapse deterioration is a hallmark of Alzheimer’s disease (AD), representing a key pathophysiological event that underlies cognitive impairment. Progressive synapse loss is strongly correlated with both the accumulation of pathological tau protein in the brain and with cognitive decline in AD [1–3]. Pathogenic tau causes synaptotoxicity that manifests through the dysregulation of signaling pathways and mechanisms at both presynaptic and postsynaptic compartments across models of AD and other tauopathies [4, 5]. The process of tau-mediated synapse deterioration by which intact presynaptic and postsynaptic compartments become dysfunctional leading to the loss of whole synaptic structures is not well understood.

Modifications on tau, including phosphorylation and acetylation, cause tau to accumulate and form small soluble oligomeric aggregates [6–8]. Growing evidence points to soluble tau oligomers as the major neurotoxic species of aggregated tau in the brain that cause synapse dysfunction underlying cognitive impairment [4, 9–15]. Synapses in postmortem AD brain tissue contain elevated levels of tau oligomers compared to healthy control brains [16–19]. Moreover, cognitively resilient individuals with AD pathology have significantly lower levels of tau oligomers at synapses [20]. In AD brain, tau oligomers are detected in both presynaptic and postsynaptic compartments, signifying that tau oligomers may directly impact synapse function and propagate in the brain through transsynaptic spreading [19]. Tau oligomers can be secreted from neurons [21, 22], and levels of tau oligomers are increased in human cerebrospinal fluid of AD patients [23], linking extracellular tau oligomers to AD-related pathogenesis. Neurons internalize extracellular tau oligomers, which trigger the propagation of endogenous tau pathology and toxicity [24–27]. Consequently, the uptake of extracellular tau oligomers in neurons could have both immediate and long-term effects on synapse function and neuronal viability.

Long-term potentiation (LTP) at synapses is a key correlate of memory encoding in the hippocampus [28], and LTP is inhibited by tau oligomers in mouse models [24, 29–34]. Even a brief 20-min administration of extracellular tau oligomers impairs LTP at synapses of the hippocampus [29], suggesting that tau oligomers immediately inhibit a mechanism required for LTP expression at synapses. Moreover, synapses acutely exposed to oligomeric tau exhibit enhanced long-term depression (LTD) [34]. Longer-term effects of tau oligomer exposure include reduced levels of presynaptic proteins in hippocampus [13] and loss of dendritic spines [35, 36]. In human induced pluripotent stem cell (iPSC)-derived neuron cultures, the number of presynaptic compartments containing Synapsin or Synaptophysin were reduced two weeks after oligomeric tau exposure [27]. While synapses containing tau oligomers can be eliminated by microglia or astrocytes [18], the functional weakening of individual synapses could also lead to their retraction [37, 38]. Indeed, synaptic mechanisms that promote long-term depression (LTD) and drive synapse elimination are also upregulated in AD models [39–41]. How acute tau oligomer-mediated synapse dysfunction may lead to the weakening and gradual deterioration of synapses is largely unknown.

In this study, we investigated how a brief tau oligomer exposure leads to the functional deterioration of glutamatergic synapses in human iPSC-derived neurons. Using a postsynaptic proteomic mapping approach, we discovered dynamic time-dependent changes in the postsynaptic proteome following oligomeric tau exposure that link disease-related processes and the dysregulation of synapse strength. Then by monitoring bipartite synapse physiology, we delineated the progressive and coinciding postsynaptic and presynaptic dysfunction caused by tau oligomers over time. Our findings reveal the series of pathophysiological events underlying the progression of tau oligomer-induced synaptotoxicity, from the immediate impact on synaptic plasticity to the eventual degeneration of synapses.

## METHODS

### Protein purification and generation of tau oligomers

C-terminal 6x His-tagged recombinant tau 2N4R in pET21-b was transformed into *E. coli* (Rosetta 2, DE3) cells and plated on LB + ampicillin agar plates. A single colony was incubated overnight and subcultured in LB broth at 37°C for 2 hours. Protein expression was induced with 0.5 mM IPTG at 37°C for 2 hours. Cells were pelleted and lysed by freeze thaw method in PBS with 2% Triton and protease inhibitor cocktail (Sigma). Streptomycin sulfate (Thermo) was added to precipitate the DNA. The lysate was boiled for 10 min and incubated on ice for 5 min before spinning down at 18,000 x g for 15 min at 4°C. The supernatant was loaded onto TALON cobalt beads (Takara Bio USA, Inc.) for affinity chromatography purification of His-tagged tau protein. The column was washed with 10 column volumes of binding buffer (50 mM Tris-HCl pH 7.4 plus 0.3 M NaCl) and 10 column volumes of binding buffer containing 5 mM Imidazole. The protein was eluted with binding buffer containing 150 mM Imidazole. Eluted protein was concentrated with Amicon Ultra Centrifugal Devices (Millipore) and loaded onto size exclusion chromatography (HiLoad Superdex 200 pg) equilibrated with PBS pH 7.4 plus 5 mM DTT and 5 mM EDTA to separate tau from smaller truncated tau. Fractions were run on SDS-PAGE for analysis. Pooled tau fractions were buffer exchanged in PBS pH 7.4 using PD10 desalting column (Sigma) to remove DTT and treated with 1 mM H_2_O_2_ overnight at room temperature to form tau oligomers through disulfide bond formation. Tau oligomers were buffer exchanged in PBS pH 7.4 using a PD10 desalting column and saved in -80°C. Pooled tau monomers were buffer exchanged in PBS pH 7.4 to remove DTT using a PD10 desalting column and saved in -80°C. Presence of tau oligomers were confirmed through SDS-PAGE and western blot. Tau-FITC monomer (rPeptide) was oligomerized using the same procedures with H_2_O_2_.

### Cell Cultures

Cell cultures were maintained in 37°C incubator with 5% CO_2_. Human iPSCs (WTC11) engineered with inducible neurogenin 2 (NGN2), described previously [42], were used to generate human neurons. Karyotyping was used to establish intact chromosomes in the iPSCs. Human iPSCs were grown in E8 media on cell culture plates coated with Matrigel (Corning). To passage confluent iPSC cultures, the cells were dissociated using Accutase (STEMCELL Technologies, Inc.) and plated in E8 media containing 10 μM ROCK inhibitor at low density (< 30% confluence). ROCK inhibitor was removed during media changes on higher density iPSC cultures. For differentiation, the human iPSCs were plated at a density of 2.0 x10^6^ cells per well onto matrigel-coated (Corning) 6-well plate in pre-differentiation Knockout DMEM/F-12 media containing 2 μg/mL doxycycline (Sigma), N2 supplement (Gibco), MEM non-essential amino acid (Gibco), 10 ng/mL brain-derived neurotrophic factor (BDNF) (Peprotech), 10 ng/mL neurotrophin-3 (NT3) (Peprotech) and 10 μM ROCK inhibitor (Y-27632, Cayman chemicals). The media was changed the following day without ROCK inhibitor and subsequently for another day. For APEX2-PSD-95 experiments, lentivirus was added on the 1^st^ day of media change without the ROCK inhibitor. After a total of 3 days in pre-differentiation media, on Day 0, the precursor cells were dissociated using Accutase and plated onto matrigel-coated coverslips or tissue culture plates with Neurobasal-A media (Gibco) containing B27 supplement (Gibco), GlutaMAX (Gibco), 10 ng/mL BDNF, 10 ng/mL NT3, and 2 μg/mL doxycycline. Within 24 h of plating the human neurons, cultured rat astrocytes (ScienCell) were added. On Day 5-6, half of the media was replaced with the Neurobasal-A media with supplements without doxycycline and with added cytosin β-D-arabinofuranoside (Ara-C) to inhibit astrocyte growth. On Day 10-11, half of the media was removed and replenished with MEM media containing 27.7 mM glucose (Sigma), 2.4 mM NaHCO_3_ (Sigma), B-27 supplement, 0.5 mM L-glutamine (Gibco), Ara-C and 5% Fetal Bovine Serum (FBS) (Gibco). Human neurons were maintained and allowed to mature by replacing one third of the supplemented MEM culture media with the same fresh media once a week for 6-8 weeks before experiments. For immunocytochemistry, human neurons were plated at a density of 200,000 cells/well with astrocytes at a density of 40,000 cells/well in 24-well plate. For mass spectrometry experiments, 8 x 10^6^ neurons were plated with 500,000 rat astrocytes per 10 cm tissue culture plate.

For HEK293 cells (Invitrogen) used for shRNA experiments, cells were maintained in DMEM media (Gibco) containing 10% FBS (Gemini), 200 mM L-glutamine, 10 mM MEM non-essential amino acids and 100 mM Sodium Pyruvate (Sigma). Lipofectamine 2000 Transfection Reagent (Invitrogen) was used for transfecting HEK293 cells with plasmids.

### SDS-PAGE and Western blot

To analyze purified tau monomers and oligomers, the protein concentrations were quantified using a Bradford Assay (Bio-Rad) and 1 μg of each protein samples were loaded twice to run in 2x 4-12% gradient SDS-PAGE gel (Invitrogen). One gel was stained with the Coomassie dye R-250 (Imperial Protein Stain, ThermoFisher) for 1 hour at room temperature and destained with water overnight before imaging with ChemiDoc MP imaging system (Bio-Rad). The proteins from another gel were transferred to a nitrocellulose membrane (GE Healthcare), incubated with blocking solution containing 5% nonfat dry milk in TBST for 1 hour at room temperature and incubated overnight with primary antibody diluted in blocking solution. The membrane was washed 3 times with TBST for 10 min at room temperature and incubated with HRP-conjugated secondary antibody diluted in TBST with 2% nonfat dry milk for 1 hour at room temperature. Then following 3 times of TBST washes, the membrane was incubated with enhanced chemiluminescence (ECL) reagent (Pierce) for 1 min at room temperature and the chemiluminescence from immunolabeled proteins was exposed on film (Thermo Scientific) and scanned. The bands from images were quantified using Fiji (ImageJ).

For HEK293 and human neuron samples, cells were scraped off from plates with RIPA buffer containing 50 mM Tris, pH 7.5, 150 mM NaCl, 0.5% Nonidet P-40, 1 mM EDTA, 0.5% sodium deoxycholate, 0.1% SDS, 1 mM phenylmethyl sulfonyl fluoride, protease inhibitor cocktail (Sigma), phosphatase inhibitors 2 and 3 (Sigma), then sonicated 10 times with 1 sec pulses. Lysates were incubated on ice for 30 min before spinning at 18,000 g at 4°C for 20 min. The supernatant was separated from the pellet and the protein concentrations were measured and further analyzed using western blot as described above.

### Exposure of human neurons to recombinant tau oligomers

At 6-8 weeks old, human neurons were treated with filter-sterilized PBS (untreated control), 6 μg/mL tau monomers, tau oligomers or oligomerized tau-FITC for 30 min at 37°C diluted in 1/3 of media while 2/3 of the media was saved. Cells were washed with warmed DPBS, and saved media was added back before returning the treated cells to the 37°C incubator until used for further experiments.

### Immunocytochemistry and imaging

Human neurons were plated on coverslips in 24-well plates. Coverslips were fixed in 4% paraformaldehyde for 15 min at room temperature and washed with PBS (Corning) three times for 5 min. For immunostaining Synapsin, GFAP and GluA2/3, neurons were incubated with blocking solution containing PBS, 0.3% Triton X-100 (Thermo) and 2% normal goat serum (NGS) (Jackson Immunoresearch) for 1 h at room temperature. Cells were incubated with primary antibodies diluted in blocking solution for 1 h at room temperature and washed with PBS three times for 5 min. Then cells on coverslips were incubated with secondary antibodies diluted in blocking solution for 1 h at room temperature, washed with PBS three times for 5 min before mounting with Prolong Gold (Invitrogen) on glass slides. For immunostaining MAP2, NeuN, Myosin-Va, and vGluT1, the same procedures were followed but with 0.1% Triton X-100 for blocking solution and antibody dilution. For experiment showing biotinylation with APEX2-PSD-95 and immunostaining for Flag, PSD-95 and Cy3 Streptavidin, the same procedures were also followed except with added 3% non-fat dry milk for blocking solution and antibody dilution. For GluN1 immunostaining, cold (-20°C) methanol was used to fix the cells at 20°C for 10 min. Cells were washed with PBS three times for 5 min before incubating with blocking solution containing PBS, 0.1% Triton X-100 and 10% NGS. Cells were incubated with primary antibodies diluted in PBS with 3% NGS for 2 h at room temperature, washed with PBS three times, then incubated with secondary antibodies in the same buffer. Coverslips were washed with PBS three times before being mounted on glass slides. Cells were imaged using either a Zeiss LSM 700 or Nikon AX laser scanning confocal microscope with a 63x oil objective and images were analyzed using FIJI software (ImageJ). Imaging and analyses were performed blind.

### Antibodies

Primary antibodies used for western blot and immunocytochemistry were: Tau5 (AB_80579, Abcam), Streptavin HRP (S911, Invitrogen), GAPDH (MAB374, Sigma), Rabbit FLAG (F7425, Sigma), Mouse FLAG (F3165, Sigma), PSD-95 (MA1046, ThermoFisher), GluA2/3 (AB1506, MilliporeSigma), Cy3 Streptavidin (AB_2337244, Jackson Immunoresearch), Mouse vGluT1 (MAB5502, MilliporeSigma), Guinea Pig vGluT1 (AB5905, MilliporeSigma), GluA1 (ABN241, MilliporeSigma), GluN1 (114 011, SynapticSystems), Synapsin (5297S, Cell Signaling), Rabbit MAP2 (4542S, Cell Signaling), Chicken MAP2 (NB300 213, Novus Biologicals), NeuN (AB177487, Abcam), GFAP (PA110004, Invitrogen)

Secondary antibodies used were: Anti-mouse Alexa Fluor 488 (115-545-166, Jackson Immunoresearch), Anti-guinea pig Alexa Fluor 546 (A-11074, Life Technologies), Anti-mouse Alexa Fluor 546 (A-11030, Invitrogen), Anti-rabbit Alexa Fluor 647 (111-605-144, Jackson Immunoresearch), Anti-chicken Alexa Fluor 647 (A32933, Invitrogen)

### Chemical LTP

At 6-8 weeks old, human neurons were treated with either 6 μg/ml of tau oligomers or tau monomers for 30 min at 37°C. Chemical LTP (cLTP) on human neurons were performed after the treatment. Neurons were first washed once with warmed (37°C) extracellular solution (ECS) containing 125 nM NaCl, 5 mM KCl, 25 mM HEPES, 1 mM NaH_2_PO4, 11 mM Glucose, and 2.5 mM CaCl_2_ (all from Sigma Aldrich) with 0.5 mM tetrodotoxin (Abcam) and 1 mM strychnine (Sigma-Aldrich) at pH 7.4. Then cells were either treated with ECS alone for the unstimulated control or with 300 μM Glycine (Sigma-Aldrich) for 15 min at 37°C to induce cLTP. Then cells were washed with ECS again, incubated at 37°C for 10 min and immunolabeled for anti-GluA1 diluted in ECS for 20 min at 37°C to monitor insertion of surface AMPARs. Cells were then washed once with ECS then fixed with 4% paraformaldehyde (PFA) for 15 min. Coverslips were washed with PBS three times for 5 min before incubating with blocking solution of PBS with 2% NGS and 0.1% Triton X-100 for 1 h. Cells were immunolabeled with anti-vGluT1 antibody in blocking solution for 1 h at room temperature and washed with PBS three times for 5 min. Then cells were labeled with anti-mouse Alexa Fluor 488 and anti-rabbit Alexa Fluor 647 secondary antibodies diluted in blocking solution for 1 h at room temperature, washed with PBS three times for 5 min and mounted. Images were acquired on a Zeiss LSM 700 and GluA1 integrated intensity colocalized with vGluT1 was measured using Fiji software (ImageJ). For the 14-day timepoint, human neurons were treated with 6 μg/ml of tau oligomers or mock-treated with PBS for 30 min at 37°C and washed with DPBS. The treated cells were incubated back with old Neurobasal media for 14 days at 37°C before performing chemical LTP. Imaging and analyses were performed blind.

### Electrophysiology

Whole-cell voltage clamp recordings on human neurons were performed at 30°C in an external solution containing (in mM) 140 NaCl, 5 KCl, 10 glucose, 10 HEPES, 2 MgCl_2_ and 2.5 CaCl_2_, pH 7.4. The 3-7 MΩ patch pipette was filled with an internal solution containing (in mM): 120 CsCl, 10 HEPES, 10 EGTA, 5 NaCl, 1 MgCl_2_, 4 Mg-ATP, 0.3 Na_3_-GTP, 10 QX-314, pH 7.2. Cells were held at -60 mV and extracellular field stimulation was applied using a concentric bipolar electrode positioned 50-100 μm from soma. A current injection of 15 μA was used to evoke excitatory postsynaptic currents (EPSCs). To measure the paired-pulse ratio (PPR), two evoked EPSCs with 20 ms interstimulus interval were recorded from neurons in external solution containing 1 mM CaCl_2_. The average of 3 pairs of evoked EPSCs (EPSC amplitude of pulse 2/EPSC amplitude of pulse 1) per cell was calculated as the PPR and used for analysis. WinLTP software (version 1.11b, University of Bristol) and a Multiclamp 700B amplifier (Molecular Devices) were used for recording acquisition. Recordings were acquired with a 10 kHz sampling rate, and a low-pass 2 kHz Bessel filter was applied post-acquisition before analyses were performed. The amplitudes of postsynaptic responses were analyzed using Clampfit (Molecular Devices). The stimulus artifact was removed from representative trace examples.

### Transmission electron microscopy (TEM)

Human neurons were plated on a 6-well plate with a density of 800,000 cells/well. When the neurons were 6 weeks old, 2 wells were exposed once to 6 μg/ml of tau oligomers for 30 min for the 14 days tau oligomer exposure group. Another 2 wells were treated with PBS as the vehicle control group. When the neurons were 7 weeks old, 2 more wells were exposed once to 6 μg/ml of tau oligomers for 30 min for the 7 days tau oligomer exposure group. When the neurons were all 8 weeks old, they were washed with wash buffer (0.1 M sodium cacodylate and 2 mM CaCl_2_, pH 7.4) before fixing with room temperature 2.5 % glutaraldehyde, 4% PFA in 0.1M sodium cacodylate buffer containing 0.02% picric acid. The next day, fixed cells were washed with chilled wash buffer four times. The samples were prepared by standard processing and embedding for TEM [43] at the Microscopy Core Facility of Weill Cornell Medicine. The embedded samples were stained with Toluidine Blue then ultrathin sections were cut and collected on copper grids for TEM imaging following standard operating conditions and viewed on a JEOL JEM1400 (JEOL Ltd, USA, Peabody, MA). Images were captured using a Veleta 2K x 2K CCD camera using RADIUS software (EMSIS, Muenster, Germany).

### Generation of APEX2-PSD-95 construct and lentivirus

A TRC2-pLKO puro plasmid with human PSD-95 shRNA (Sigma-Aldrich, TRCN0000235627), comprised of the following sequence: ‘ACGAGAGTGGTCAAGGTTAAACTCGAGTTTAACCTTGACCACTCTCGT’, was co-expressed with human PSD-95 (pcDNA3.1-C-(k)DYK) in HEK293 cells to test knockdown. To make the APEX2-PSD-95 construct, APEX2 [44] and the flag tag sequence was fused to the C-terminus of rat PSD-95 cDNA [45] (Addgene plasmid #74036). Three silent mutations were introduced into the rat PSD-95 cDNA, corresponding to residues E498, S500, and L502, to make it resistant to the PSD-95 shRNA. The human PSD-95 shRNA and APEX2-PSD-95 were inserted downstream of a U6 promotor and human ubiquitin C promoter, respectively, in an FUGW2 lentiviral plasmid (from Dr. Li Gan). The APEX2-PSD-95 construct was synthesized by GenScript and sequence was confirmed by Sanger sequencing. Lentivirus to express APEX2-PSD-95 in human iPSC-derived neurons was prepared in HEK293 cells. HEK293 cells were split and plated on 15 cm cell culture plates. The following day, a calcium phosphate transfection method was used to transfect HEK293 cells (50-60% confluent) with the APEX2-PSD-95 construct along with Δ8.9 and VSV-G packaging plasmids (from Dr. Li Gan). Media was replaced the next day and lentivirus was harvested 24 and 48 hours later. The collected virus was filtered with 0.45 μm vacuum filter (Fisher Scientific), purified with sucrose gradient ultracentrifugation and resuspended in sterile PBS. The virus titer was quantified using p24 Rapid Titer Kit (Takara Bio USA, Inc.). Additional lentivirus was purchased from VectorBuilder.

### APEX proximity labeling

After treatment of cells with vehicle or tau oligomers, cells were washed with warmed (37°C) ACSF containing (in mM): 140 NaCl, 5 KCl, 2.5 CaCl_2_, 2 MgCl_2_, 10 HEPES, 10 glucose at pH 7.4. Then cells were incubated with 500 μM biotin-phenol (Adipogen) diluted in ACSF for 30 min at 37°C cell culture incubator. Cells were treated with 1 mM hydrogen peroxide (H_2_O_2_) for 1 min then immediately quenched with cold quenching solution containing PBS with 10 mM sodium azide, 10 mM sodium ascorbate, and 5 mM Trolox twice (all reagents from Sigma). After removing the quenching solution, cells were either fixed for immunostaining or lysed for pull-down. The cells were lysed in buffer containing 50 mM Tris-Cl pH 7.4, 500 mM NaCl, 0.5% NP40, 1 mM EDTA, 1 mM DTT, 1 protease inhibitor tablet (Pierce), phosphatase inhibitors 2 & 3, 10 mM sodium azide, 10 mM sodium ascorbate, and 5 mM Trolox. Lysates were then sonicated with a sonicator (SONICS) at output level 3, 30% duty cycle for 30 pulses while submerged in ice then centrifuged at 16,500 g for 10 min at 4°C. Supernatant from samples were stored in low-protein binding tubes and stored in -80°C.

### Immunoprecipitation of biotinylated postsynaptic proteins

Streptavidin-coated magnetic beads (Pierce #88817) were washed twice with lysis buffer containing 50 mM Tris-HCl, pH 7.4, 150 mM NaCl, 0.5% NP-40, 1 mM EDTA, 1 mM DTT. Protein concentration from human neuron lysates were measured using a Bradford assay (Bio-Rad #5000006) and 2 mg of each sample were incubated with 200 ml of washed Streptavidin-coated magnetic beads overnight at 4°C. Beads were washed four times with wash buffer containing 50 mM Tris-HCl, pH 7.4, 150 mM NaCl, 0.5% NP-40, 0.5% deoxycholate, 1 mM EDTA, 1 mM DTT and then washed four times with 1 ml of 50 mM Ammonium Bicarbonate at 4°C for 8 min. Washed beads were then resuspended in 100 ml 2 M Urea dissolved in 50 mM Ammonium Bicarbonate. Samples were reduced on resin with 5 mM TCEP with orbital shaking (1200 rpm) for 30 min at 37°C. Samples were then alkylated with 20 mM iodoacetamide in the dark with orbital shaking (1200 rpm) for 30 min at room temperature and quenched with 20 mM DTT with orbital shaking (1200 rpm) for 15 min at room temperature. Solution from each sample was removed and beads were resuspended in 100 ml of 1 M Urea in 50 mM Ammonium Bicarbonate.

### Protein Digestion and Desalting

Protein samples previously reduced with tris(2-carboxyethyl)phosphine (TCEP), alkylated with iodoacetamide and resuspended in Urea were incubated overnight at 37 °C with sequencing grade trypsin (Promega, San Luis Obispo, CA) dissolved in 50 mM TEAB at ∼1:25 (w/w) enzyme:protein ratio. The following day, peptides were extracted from streptavidin-coated magnetic beads by using a magnet. Afterwards, samples were dried down by centrifugal evaporation, resuspended in 0.2% FA in water, and desalted with in-house C_18_ packed stage-tips. Desalted peptides were dried down by centrifugal evaporation and re-suspended in 20 µL of 0.2% FA in water. Peptides were diluted by 1 to 20 with 2% ACN, 0.1% FA in water and 1 µL of indexed Retention Time Standard (iRT, Biognosys, Schlieren, Switzerland) was added to each sample.

### Mass spectrometry analysis

Reverse-phase HPLC-MS/MS analyses were performed on a Dionex UltiMate 3000 system coupled online to an Orbitrap Exploris 480 mass spectrometer (Thermo Fisher Scientific, Bremen, Germany). The solvent system consisted of 2% ACN, 0.1% FA in water (solvent A) and 80% ACN, 0.1% FA in ACN (solvent B). Digested peptides (2 µL, 1 to 20 dilution) were loaded onto an Acclaim PepMap 100 C_18_ trap column (0.1 x 20 mm, 5 µm particle size; Thermo Fisher Scientific) over 5 min at 5 µL/min with 100% solvent A. Peptides were eluted on an Acclaim PepMap 100 C_18_ analytical column (75 µm x 50 cm, 3 µm particle size; Thermo Fisher Scientific) at 300 nL/min using the following gradient: linear from 2.5% to 24.5% of solvent B in 125 min, linear from 24.5% to 39.2% of solvent B in 40 min, up to 98% of solvent B in 1 min, and back to 2.5% of solvent B in 1 min. The column was re-equilibrated for 30 min with 2.5% of solvent B, and the total gradient length was 210 min. Each sample was acquired in data-independent acquisition (DIA) mode [46–48]. Full MS spectra were collected at 120,000 resolution (Automatic Gain Control (AGC) target: 3e6 ions, maximum injection time: 60 ms, 350-1,650 *m/z*), and MS2 spectra at 30,000 resolution (AGC target: 3e6 ions, maximum injection time: Auto, Normalized Collision Energy (NCE): 30, fixed first mass 200 *m/z*). The isolation scheme consisted of 26 variable windows covering the 350-1,650 *m/z* range with an overlap of 1 *m/z* [49].

### DIA Data Processing and Statistical Analysis

DIA data was processed in Spectronaut (version 19.4.241104.62635) using directDIA. Data extraction parameters were set as dynamic and non-linear iRT calibration with precision iRT was selected. Data was searched against the *Homo Sapiens* reference proteome with 20,411 entries (UniProtKB), accessed on 10/06/2023. Trypsin/P was set as the digestion enzyme and two missed cleavages were allowed. Cysteine carbamidomethylation was set as a fixed modification while methionine oxidation and protein N-terminus acetylation were set as dynamic modifications. Identification was performed using 1% precursor and protein q-value. Quantification was based on the peak areas of extracted ion chromatograms (XICs) of 3 – 6 MS2 fragment ions, specifically b- and y-ions and q-value sparse data filtering applied. In addition, iRT profiling was selected. Differential protein expression analysis comparing 1) oTau to NTS (control) was performed using a paired t-test, and p-values were corrected for multiple testing, using the Storey method [50]. Specifically, group wise testing corrections were applied to obtain q-values. Protein groups with at least two unique peptides, p-value < 0.05, and absolute Log_2_(fold-change) > 0.25 are significantly altered.

### Data Availability

Raw data for images and immunoblots are available upon request. Raw data and complete MS data sets have been uploaded to the Mass Spectrometry Interactive Virtual Environment (MassIVE) repository, developed by the Center for Computational Mass Spectrometry at the University of California San Diego, and can be downloaded using the following link: https://massive.ucsd.edu/ProteoSAFe/dataset.jsp?task=41cbdbf0c4784f3795d0e745a8ad37e5 (MassIVE ID number: MSV000097844; ProteomeXchange ID: PXD063816).

### Enrichment analyses

Proteins identified from both mass spectrometry experiments were analyzed with Synaptic Gene Ontologies (SynGO) cellular component [51]. Significantly downregulated proteins in tau oligomer treated neurons 1 hour after treatment were analyzed using Gene Set Enrichment Analysis (GSEA) [52, 53]. Significantly upregulated proteins were analyzed for reactome pathways using STRINGdb [54] and modified for visualization with Cytoscape version 3.10.2 [55]. Graphs from enrichment analyses were created using R programming showing p-values and fold changes of enriched proteins. Illustrations, schematic and venn diagrams were created using BioRender (BioRender.com).

### Statistical Analyses

Student’s *t*-test was used to measure the statistical differences between the means of two groups while one-way ANOVA with Bonferroni *post*-hoc analyses was used to analyze the differences between means of multiple groups.

## RESULTS

### Acute tau oligomer exposure leads to impaired LTP and immediate changes in the postsynaptic proteome

To investigate tau oligomer-mediated synaptotoxicity in a human neuron model, we used human iPSCs from a healthy individual (WTC11) with genetic integration of a doxycycline inducible NGN2 transgene. NGN2 expression was used to initiate the differentiation of the iPSCs into glutamatergic neurons which were then cultured for ≥ 6 weeks until experiments were performed when the neurons established functionally mature synapses [42]. Since tau oligomers immediately block LTP in mouse models [29], we first evaluated the acute effect of tau oligomers on LTP in human neurons. Human recombinant tau (2N4R) monomers were purified and treated for 20 hours with hydrogen peroxide to generate tau oligomers [29]. Tau oligomer formation was confirmed by Coomassie blue staining and western blot (Suppl. Fig. S1A, S1B). Human neurons were treated with either tau oligomers or tau monomers starting at 30 min before the induction of chemical LTP (cLTP), which involves the activation of synaptic NMDA receptors (NMDAR) to drive the postsynaptic insertion of additional AMPA receptors (AMPARs) thereby enhancing synapse strength [56, 57]. To monitor postsynaptic AMPAR trafficking after cLTP, we performed surface immunolabeling of GluA1 on live neurons followed by fixation, membrane permeabilization and immunostaining of a presynaptic marker, vesicular glutamate transporter 1 (vGluT1), to label synapses (Fig. 1A). Immunolabeling of neurons treated with tau monomers revealed a significant increase in postsynaptic GluA1-containing AMPARs that colocalized with vGluT1 at synapses after cLTP induction compared to unstimulated control neurons (Fig. 1B). The cLTP-dependent increase in postsynaptic GluA1 was blocked in the neurons treated with tau oligomers (Fig. 1B), which is consistent with the acute inhibition of LTP expression by tau oligomers reported in mouse hippocampus [29].

**Figure 1.**
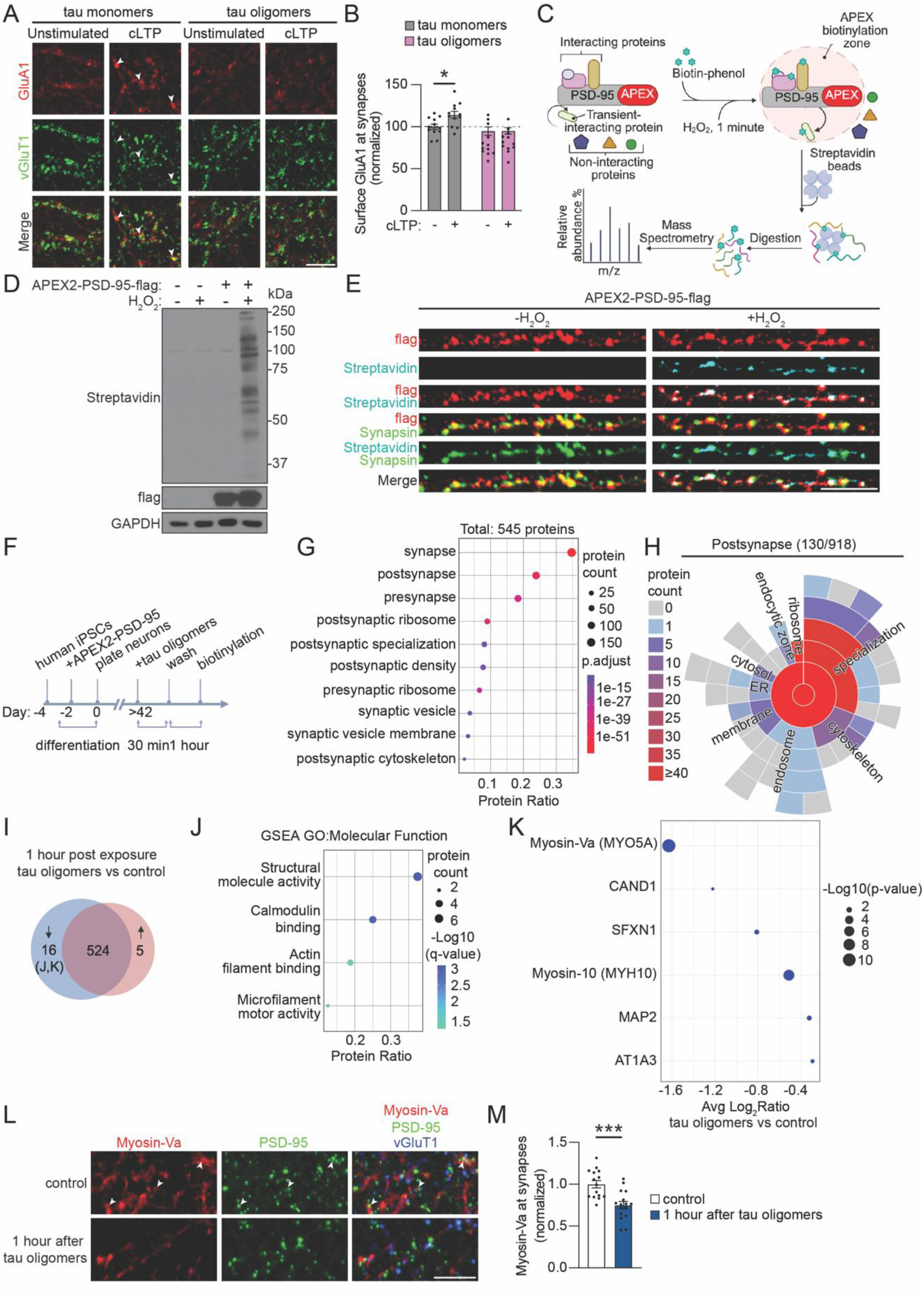
Acute tau oligomer exposure rapidly blocks LTP-induced AMPAR trafficking and downregulates myosins at postsynaptic sites in human neurons. **A** Representative confocal images of immunolabeling of surface GluA1-containing AMPARs (red) and vGluT1 (green), a synaptic marker, in human neurons. Neurons were treated with tau monomers or oligomers for 30 min and then they were either unstimulated or stimulated with chemical induction of LTP (cLTP). The neurons were fixed at approximately 30 min after cLTP induction. Scale bar: 5 μm. **B** Quantification of the intensity of surface GluA1 at synapses normalized to unstimulated neurons treated with tau monomers (n = 12 neurons/group; * p < 0.05 Unpaired Student’s *t*-test). **C** Workflow of protocol to biotinylate postsynaptic proteins in close proximity with APEX2-PSD-95 in human neurons, to pulldown the biotinylated proteins using streptavidin beads and to identify the biotinylated peptides using mass spectrometry. **D** Immunoblot of flag-tagged APEX2-PSD-95, biotinylated proteins (streptavidin), and GAPDH loading control from human neurons with or without APEX2-PSD-5 expression and with or without H_2_O_2_ treatment for 1 min. **E** Human neurons expressing APEX2-PSD-95 incubated with biotin-phenol for 30 min and stimulated with H_2_O_2_ for 1 min. Cells were fixed and immunostained for APEX2-PSD-95 (flag, red), biotinylated proteins (Streptavidin, blue) and a synaptic marker (Synapsin, green). Scale bar: 5 μm. **F** Workflow of treatment of human neurons with tau oligomers for proximity labeling of postsynaptic proteins. **G** SynGO cellular component analyses of 545 biotinylated proteins identified by mass spectrometry from neurons that were untreated or exposed to tau oligomers. Size of circles denotes protein count, colors denote p-values and protein ratio denotes protein count over total number of proteins. **H** SynGO analysis showed 130 total annotated postsynaptic proteins including the proteins classified in child terms. **I** Venn diagram analysis of the 545 biotinylated proteins detected in untreated neurons or in neurons 1 hour after tau oligomer exposure highlighting the significantly upregulated (red) and downregulated (blue) proteins in the tau oligomer-treated neurons compared to untreated neurons (n = 5-6 cultures/group; p < 0.05 with absolute average log_2_ratio > 0.25). **J** Gene Set Enrichment Analysis on significantly downregulated proteins in tau oligomer-treated neurons showing GO molecular function. **K** Analyses of the biotinylated proteins that were significantly downregulated in tau oligomer-exposed neurons. These proteins were also reduced in postmortem AD brain tissue [68]. **L** Representative confocal images of immunostaining of Myosin-Va (red), PSD-95 (green) and vGluT1 (blue) in human neurons treat with tau oligomers for 30 min and fixed 1 hour later. Scale bar: 5 μm. **M** Quantification of the postsynaptic Myosin-Va immunolabeling intensity that colocalized with postsynaptic (PSD-95) and presynaptic (vGluT1) markers in human neurons treated with tau oligomers for 30 min and fixed after 1 hour (n = 15-16 images/group; ***, p < 0.001, Unpaired Student’s *t*-test).

We designed an approach to monitor the temporal changes in the postsynaptic proteome dynamics after tau oligomer treatment. Proximity labeling by ascorbate peroxidase (APEX2) [44] is a powerful tool used to map proteomes of different cellular compartments [58–61]. We engineered a proximity labeling system that is spatially enriched in the postsynaptic compartment by fusing PSD-95 with APEX2 containing a C-terminal FLAG tag (Suppl. Fig. S1C). We used a replacement strategy to minimize the effect of PSD-95 overexpression on synapse function [62], by including a short hairpin RNA (shRNA) to knockdown endogenous PSD-95 in human neurons. Silent mutations were introduced into the APEX2-PSD-95 sequence to make it resistant to the shRNA. Knockdown of PSD-95 by shRNA was confirmed in HEK293 cells (Suppl. Fig. S1D, S1E). The replacement strategy of PSD-95 was confirmed in human neurons where expression of APEX2-PSD-95 together with PSD-95 shRNA did not significantly change the intensity of PSD-95 immunolabeling that colocalized with Synapsin (Suppl. Fig. S1F, S1G).

Human iPSC-derived neurons were infected with lentivirus for expression of the APEX2-PSD-95 construct with PSD-95 shRNA after initiating differentiation. At 6 weeks of age, the APEX2-PSD-95 infected neurons were treated for 30 min with biotin-phenol followed by stimulation with H_2_O_2_ for 1 min to activate APEX2 enzymatic activity for biotinylation of postsynaptic proteins in close proximity to APEX2-PSD-95 (Fig. 1C). First, we confirmed that the biotinylation of proteins by APEX2-PSD-95 was dependent on the brief 1 min exposure to H_2_O_2_ (Fig. 1D). Human neurons expressing APEX2-PSD-95 were also fixed with or without the 1 min H_2_O_2_ exposure and stained with an anti-flag antibody to mark APEX2-PSD-95 localization and a fluorescently conjugated streptavidin to label biotinylated proteins. Strikingly, biotinylated proteins marked by streptavidin colocalized with APEX2-PSD-95 in puncta that overlapped with anti-Synapsin immunolabeling at synapses (Fig. 1E), supporting the postsynaptic enrichment of APEX2-PSD-95 and localized H_2_O_2_-dependent biotinylation at sites of synaptic contact.

We next sought to determine the effect of the acute 30-min oligomeric tau exposure on postsynaptic proteome dynamics with APEX2-PSD-95 proximity labeling in human neurons. The postsynaptic proteome was first mapped 1 hour after oligomeric tau exposure (Fig. 1F), corresponding to approximately when GluA1 trafficking was impaired after cLTP induction (Fig. 1B). We identified a total of 545 biotinylated proteins across both tau oligomer-treated and vehicle-treated control human neurons (Suppl. table S1). Synaptic Gene Ontologies (SynGO) cellular component analyses of the biotinylated proteins yielded significant enrichments in the postsynapse including the postsynaptic density, postsynaptic ribosomes and the postsynaptic cytoskeleton (Fig. 1G, Suppl. table S1). Presynapse was also a significantly enriched ontology term (Fig. 1G, Suppl. table S1), but most of the biotinylated proteins classified in the presynapse ontology term (70%) were also postsynaptic proteins (Suppl. Fig. S1H), including ribosomal proteins, chaperones and cytoskeletal proteins. A total of 130 biotinylated proteins were classified in the postsynapse by SynGO analyses (Fig. 1H, Suppl. table S1). Moreover, 204 of the biotinylated proteins were also detected in a proteomics study on the postsynaptic density fraction extracted from human brain tissue [63] (Suppl. Fig. S1I, Suppl. table S1). We found that 16 biotinylated proteins were significantly downregulated in neurons 1 hour after oligomeric tau exposure compared to vehicle-treated control neurons, and only 5 biotinylated proteins were significantly upregulated (Avg Log_2_Ratio of > 0.25, p < 0.05) (Fig. 1I, Suppl. table S1). Gene Set Enrichment Analysis (GSEA) of molecular function on the downregulated biotinylated proteins revealed features of structural molecule activity and actin filament binding (Fig. 1J, Suppl. table S1). Biotinylated Myosin-Va (MYO5A) was the most downregulated after tau oligomer exposure (Fig. 1K), suggesting a tau-mediated decrease in postsynaptic Myosin-Va levels. Myosin-Va is an actin-binding motor protein that regulates the activity-dependent trafficking of GluA1-containing AMPARs into spines [64] and the transport of surface NMDARs in hippocampal neurons [65]. Myosin-10 (MYH10) is another actin-binding motor protein with significantly reduced biotinylation by APEX2-PSD-95 after tau oligomer exposure (Fig. 1K). Myosin-10, also known as non-muscle myosin IIb, has many functions in the neurons including supporting LTP and synapse formation [66, 67]. We then compared the downregulated biotinylated proteins with a meta-analysis of postmortem brain tissues and found that 6 proteins, including Myosin-Va and Myosin-10, were decreased in AD brains compared to non-demented and asymptomatic controls [68–78] (Fig. 1K, Suppl. table S1). We further probed postsynaptic Myosin-Va levels 1 hour after tau oligomer exposure by performing immunostaining for Myosin-Va, PSD-95, and vGluT1 (Fig. 1L). Synaptic Myosin-Va immunostaining was significantly reduced 1 hour after tau oligomer exposure (Fig. 1M), confirming the postsynaptic downregulation of Myosin-Va detected by proteomics analyses. Our results show that LTP impairment and postsynaptic proteome changes, including the downregulation of postsynaptic actin-based motor proteins, are early events during pathogenesis in human neurons that emerge within 1 hour of tau oligomer exposure.

### Mapping the subsequent changes in postsynaptic proteome dynamics following acute tau oligomer exposure

To map the changes in the postsynaptic proteome over a longer period of time, we performed APEX2-PSD-95 proximity labeling 24 hours after the 30-min tau oligomer exposure (Fig. 2A). Mass spectrometry analyses identified 1,436 biotinylated proteins in both tau oligomer-treated and vehicle-treated control human neurons with major cellular component classifications in the postsynapse and the postsynaptic specialization (Fig. 2B, Suppl. table S2). Of the biotinylated proteins classified in the presynapse, 54% were also in the postsynapse gene ontology (Suppl. Fig. S2A, Suppl. table S2), representing proteins localized in both pre- and postsynaptic compartments. The 300 biotinylated proteins categorized by SynGO analyses of the postsynapse cellular component revealed an enrichment in components of the postsynaptic density (Fig. 2C, Suppl. table S2). Moreover, 463 of the biotinylated proteins overlapped with the postsynaptic density proteome in human brain tissue (Suppl. Fig. S2B, Suppl. table S2). The biotinylated proteins included ion channels (the GluA4 AMPAR subunit and the Kv1.2 potassium voltage-gated channel), transmembrane proteins (neuroligin 2/3/4, EphB1, EphB2, and the low-density lipoprotein receptor-related protein 1), cytoskeleton-associated proteins (actin, actinin, Myosin-10, the Arp2/3 complex, drebrin, tubulin, kinesin, tau, and Map2), as well as signaling molecules (CaMKIIα, CaMKIIβ, GSK3β, β-catenin), postsynaptic scaffolding proteins (PSD-95, Shank3) and ribosomal proteins (Fig. 2D, Suppl. table S2). Importantly, 86% of the proteins biotinylated by APEX2-PSD-95 1 hour after tau oligomer exposure were also detected in the analyses of biotinylated proteins at the 24-hour time point (Fig. 2D, Suppl. Fig. S2C, Suppl. table S2).

**Figure 2.**
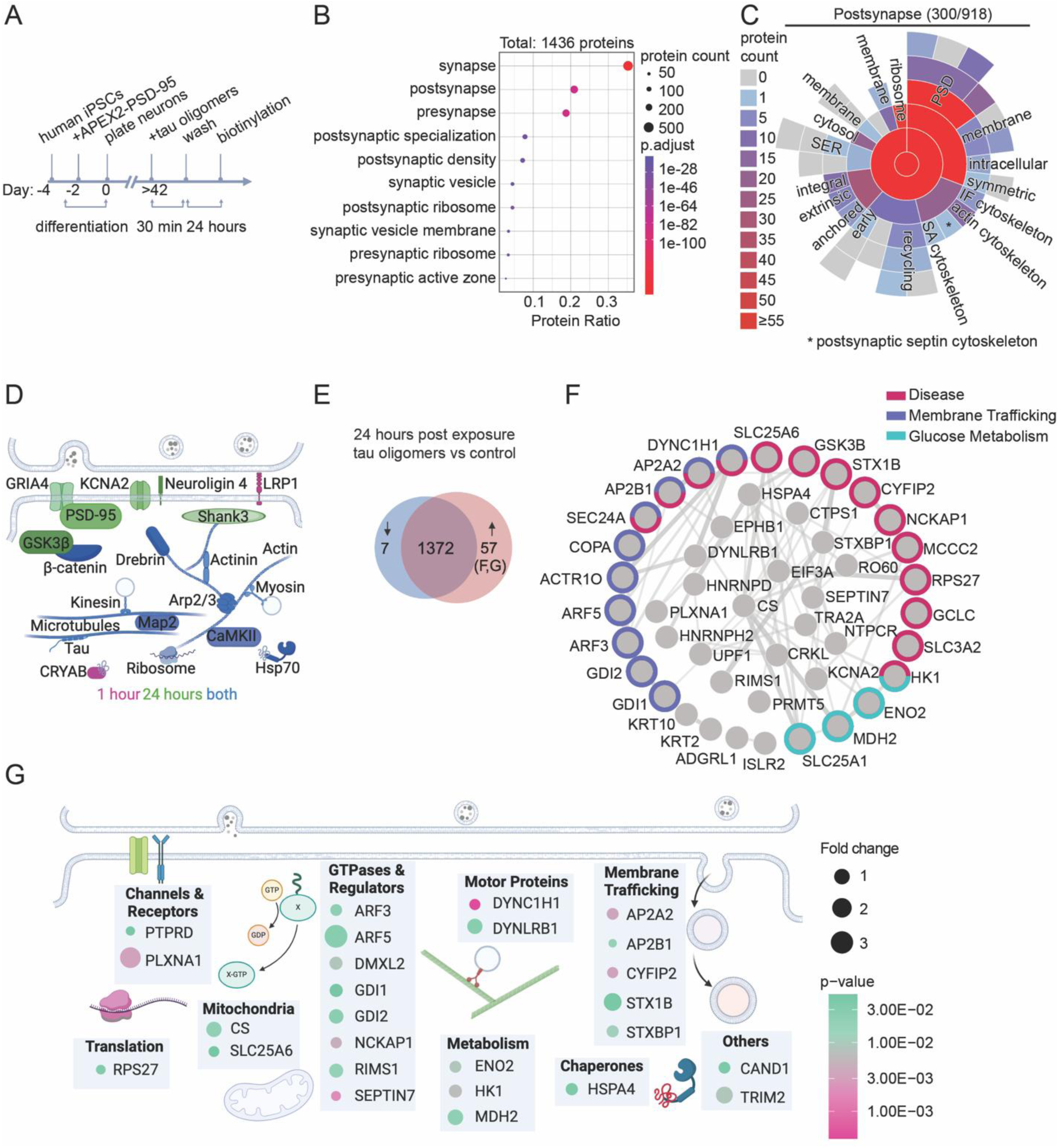
Mapping the longer-term postsynaptic proteome dynamics following acute tau oligomer exposure shows an upregulation of postsynaptic disease-related proteins. **A** Workflow of treatment of human neurons with tau oligomers followed by proximity labeling of postsynaptic proteins 24 hours later. **B** SynGO cellular component analyses of 1436 biotinylated proteins identified by mass spectrometry from untreated and tau oligomer-treated human neurons. Size of circles denotes protein count, colors denote p-values and protein ratio denotes protein count over total number of proteins. **C** SynGO analysis showed 300 annotated postsynaptic proteins including the proteins classified in child terms. **D** Schematic diagram of key biotinylated postsynaptic proteins identified in human neurons. **E** Venn diagram of the 1,436 biotinylated proteins detected from untreated neurons and tau oligomer-treated human neurons highlighting upregulated (red) and downregulated (blue) proteins at 24 hours after tau oligomer exposure compared to untreated controls (n = 6 cultures/group; p < 0.05 with absolute average log_2_ratio > 0.25). **F** Protein interaction network based on STRING database analysis of the 57 upregulated proteins in tau oligomer-treated neurons. Single proteins not shown. Colors depict Reactome pathway categories. **G** Schematic diagram of upregulated biotinylated proteins in tau oligomer-treated neurons, which were also identified in the human postsynaptic proteome, showing postsynaptic cellular classifications. Size of circles denotes fold change and colors denote p-values.

At 24 hours after tau oligomer exposure, 57 biotinylated proteins were significantly upregulated and 7 were significantly downregulated compared to vehicle-treated control neurons (Avg Log_2_Ratio of > 0.25, p < 0.05) (Fig. 2E, Suppl. table S2). Reactome pathway analyses of the 57 upregulated biotinylated proteins revealed that many are involved in disease, membrane trafficking and glucose metabolism pathways (Fig. 2F, Suppl. table S2). The glutamate-cysteine ligase catalytic subunit (Gclc), an important rate-limiting step protein for synthesizing the antioxidant glutathione [79, 80], was one of the most upregulated proteins in tau oligomer treated neurons (Fig 2F., Suppl. table S2). Gclc expression is enhanced by the transcription factor Nrf-2 in response to oxidative stress [81, 82], which supports the upregulation of a stress response in neurons 24 hours after tau oligomer exposure. Strikingly, a significant increase in GSK3β biotinylated by APEX2-PSD-95 was detected, signifying the accumulation of GSK3β at postsynaptic sites 24 hours after tau oligomer exposure. GSK3β is a key enzyme that phosphorylates tau [83, 84], and it is implicated in the pathogenesis of neurodegenerative diseases such as AD and Parkinson’s disease [85–87]. At postsynaptic sites, GSK3β activity inhibits LTP and promotes long-term depression (LTD) [88–91]. Interestingly, the tau oligomer-exposed neurons had significantly increased levels of biotinylated subunits α2 (AP2A2) and β2-adaptin (AP2B1) of the AP-2 complex which is required for AMPAR endocytosis during LTD [92, 93], supporting the upregulated postsynaptic localization of endocytic machinery related to synaptic depression (Fig. 2F, Suppl. table S2). Increased biotinylation of mitochondrial proteins included the ADP/ATP translocase 3 (SLC25A6) and several proteins involved in glucose metabolism such as tricarboxylate transport protein (SLC25A1) [94], malate dehydrogenase (MDH2) [95], hexokinase-1 (HK1) [96], and citrate synthase (CS) [97] (Fig. 2F, Suppl. table S2). This implicates altered postsynaptic mechanisms involved in mitochondrial function and glucose metabolism, which are impaired in neurodegenerative diseases [98]. The upregulated biotinylated proteins, which were also identified in the postsynaptic density extracted from human brain tissue [63], spanned additional functional classifications including GTPases and regulators, membrane trafficking proteins, and motor proteins (Fig. 2G, Suppl. table S2). Together, our temporal mapping of postsynaptic proteome dynamics suggests that tau oligomers trigger a sequential series of postsynaptic signaling events in human neurons that could lead to both immediate effects on synapse function and to the instability of synapses over time.

### Acute tau oligomer uptake is associated with a protracted deterioration in human neurons

We next investigated the uptake and stability of the tau oligomers in human neuron cultures over time. To visualize the exogenously added tau oligomers, neurons were treated with oligomerized recombinant tau conjugated to a FITC fluorophore (tau-FITC). Human neuron cultures were incubated with tau-FITC oligomers for 30 min which was followed by washing to remove the remaining extracellular tau-FITC. Then cultures were fixed for fluorescence imaging immediately after the 30-min treatment, as well as 1 hour and 24 hours after tau-FITC oligomer treatment (Fig. 3A). Internalized tau-FITC oligomers were distributed across the cultures after the 30-min oligomer treatment, both in co-cultured rat astrocytes marked by GFAP (Suppl. Fig. S3) and in synapses marked by the co-localization of postsynaptic PSD-95 and presynaptic Synapsin immunolabeling (Fig. 3B). However, at 24 hours after tau-FITC oligomer exposure, the tau-FITC fluorescence was significantly reduced at synapses (Fig. 3C, D), indicating that the internalized tau oligomers were degraded within a short period of time.

**Figure 3.**
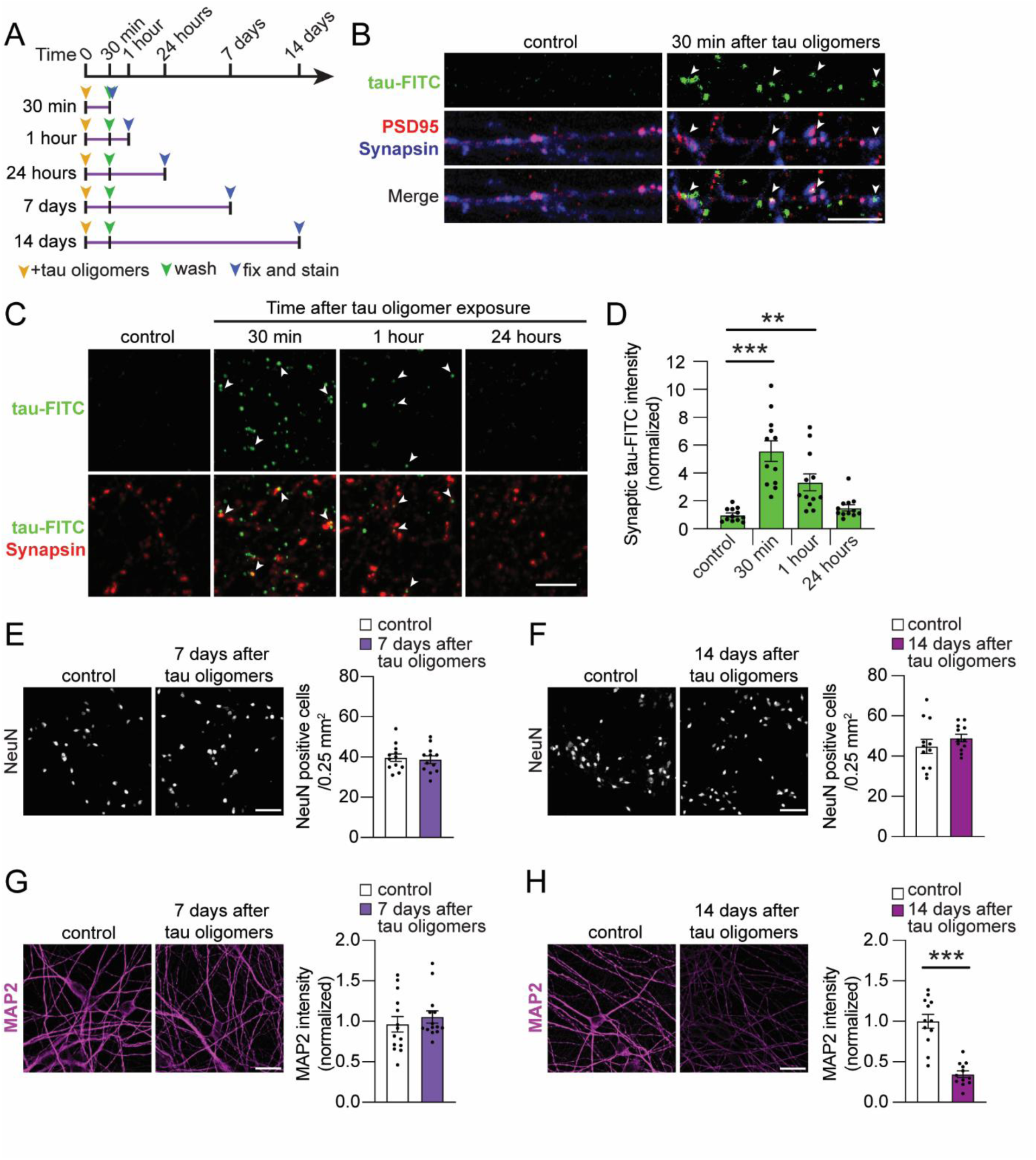
A brief tau oligomer exposure causes a prolonged degenerative effect in human neurons. **A** Workflow indicating that the 30-min treatment with tau oligomers was followed by different intervals of time (30 min, 1 hour, 24 hours, 7, or 14 days) before the fixation of human neuron cultures for experiments. **B** Confocal images of human neurons treated with oligomerized tau conjugated with Fluorescein-5-isothiocyanate (tau-FITC) (green) showed colocalization (white arrowheads) of tau-FITC with Synapsin (blue) and PSD-95 (red). Scale bar: 5 μm. **C** Representative confocal images of human neurons at 30 min, 1 hour and 24 hours after the 30 min exposure with oligomerized tau-FITC as described in **A**. Scale bar: 5 μm. **D** Quantification of the integrated intensity of tau-FITC fluorescence at synapses in human neurons that were treated with oligomerized tau-FITC for 30 min and fixed at different time points **A** (n = 12 images/group; ** p < 0.01, *** p < 0.001, one-way ANOVA, Bonferroni post hoc analyses). **E, F** Representative images of NeuN immunolabeling, as a neuronal marker, in vehicle-treated control neurons and in neurons at either (**E**) 7 or (**F**) 14 days after the 30-min exposure to tau oligomers. The density of NeuN-positive cells in the cultures was calculated and compared with or without tau oligomer exposure (n = 12 images/group; no significant difference, Unpaired Student’s *t*-test). Scale bar: 100 mm. **G, H** Representative confocal images of MAP2 immunostaining in neurons at (**G**) 7 or (**H**) 14 days after the 30-min tau oligomer treatment and in neurons treated with vehicle control. Scale bars: 20 μm. The intensity of MAP2 immunolabeling in dendrites of human neurons was quantified at (**G**) 7 days (n = 12 images/group; no significant difference, Unpaired Student’s *t*-test) and (**H**) 14 days (n = 12 images/group; *** p < 0.001, Unpaired Student’s *t*-test) post-oligomer exposure. Values are given as means ± SEM.

Exposures to extracellular pathogenic tau species can drive varying degrees of neurodegenerative phenotypes in human iPSC-derived neuron models depending on the type of toxic tau species used (oligomeric vs. fibrillar), the treatment duration and the genetic background of the human neurons used [27, 99–102]. After a short exposure of human neurons to tau oligomers for only 30 min, we did not detect a change in the density of NeuN-positive cells in the cultures at either 7 or 14 days later (Fig. 3E, F), suggesting that the brief tau oligomer exposure was not sufficient to drive human neuron loss. The dendrites of human neurons immunolabeled by MAP2 remained intact 7 days after tau oligomer exposure (Fig. 3G). However, neurons that received the acute tau oligomer exposure had significantly reduced dendritic MAP2 levels compared to control neurons after 14 days (Fig. 3H), indicating a delayed onset of cytoskeletal-related deterioration in the neurons. Overall, our results show that a brief exposure to tau oligomers is sufficient to induce protracted cytoskeletal deterioration in dendrites two weeks later.

### Progressive synapse deterioration triggered by tau oligomers

To delineate the effect of acute tau oligomer exposure on the stability of synaptic structures over time, we performed co-immunolabeling of both the presynaptic and postsynaptic compartments in the human neurons. The neurons were fixed at 2, 7 or 14 days after the acute 30-min exposure to either tau oligomers or tau monomers (Fig. 4A). Immunolabeling of Synapsin, a presynaptic cytoskeletal-associated protein that regulates synaptic vesicles, and PSD-95, the postsynaptic scaffolding protein, was performed to monitor the presynaptic and postsynaptic compartments, respectively. The co-localization of Synapsin and PSD-95 puncta represent the adjoining presynaptic and postsynaptic compartments at each synapse (Fig. 4B). We first analyzed the density of Synapsin puncta and the density of PSD-95 puncta on the human neurons separately, and then the colocalized Synapsin and PSD-95 puncta at the sites of synaptic connections were analyzed. Two days after tau oligomer exposure, the density of the Synapsin puncta, PSD-95 puncta, and the colocalized puncta were not changed compared to both vehicle-treated control neurons and neurons that were treated with tau monomers (Fig. 4B, C). Interestingly, the density of PSD-95 puncta was significantly reduced 7 days after tau oligomer exposure compared to both vehicle and tau monomer-treated groups, whereas the density of Synapsin puncta remained unchanged (Fig. 4D, 4E). This was further associated with a significant decrease in the density of synapses marked by Synapsin and PSD-95 co-localization (Fig. 4D, 4E). After one more week, at 14 days post-exposure, the density of Synapsin puncta and PSD-95 puncta were both significantly decreased on tau oligomer-treated neurons compared to vehicle-treated control and tau monomer-treated neurons, with a corresponding loss of synapses comprised of both presynaptic and postsynaptic components (Fig. 4F, 4G).

**Figure 4.**
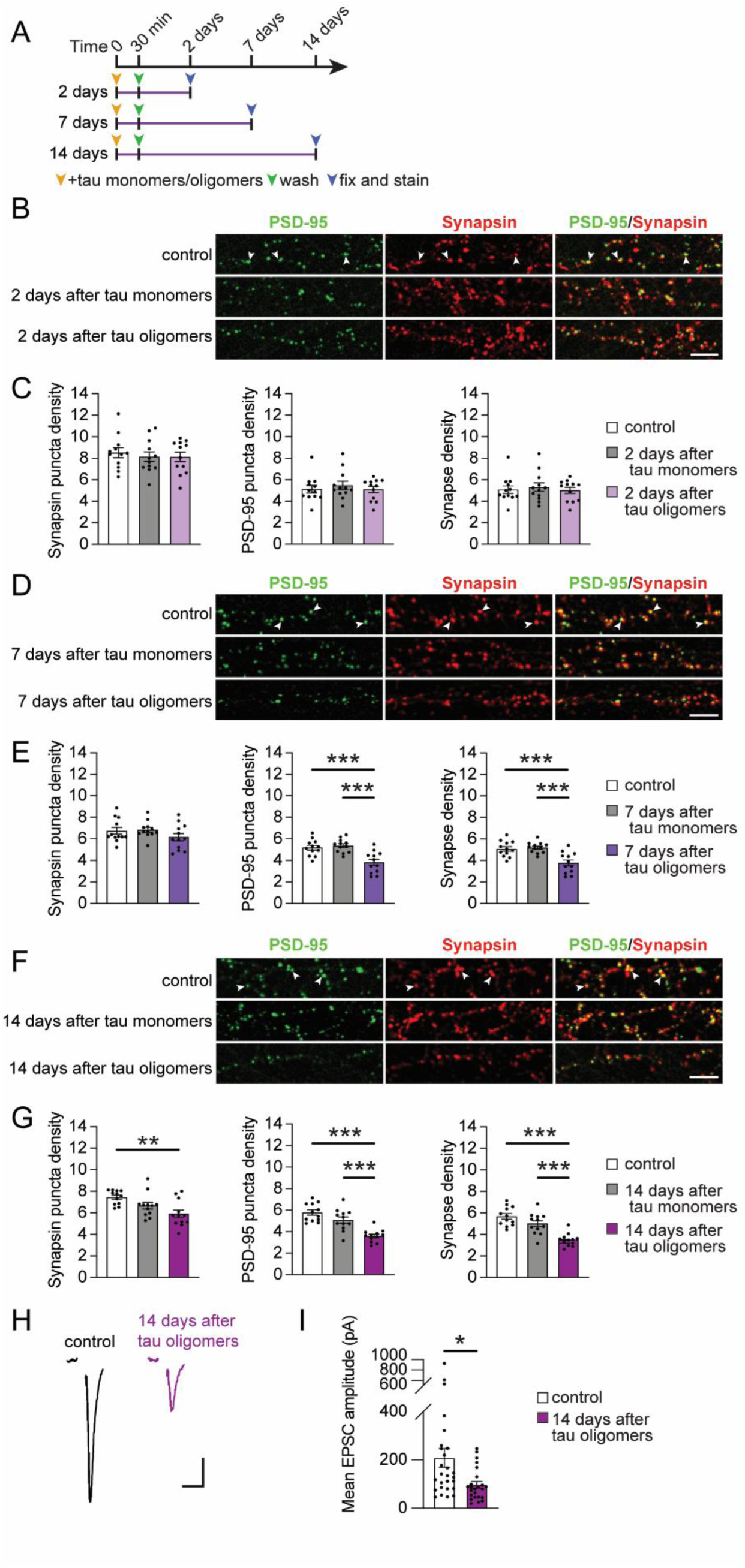
Acute tau oligomer exposure drives progressive synapse loss across postsynaptic and presynaptic compartments in human neurons. **A** Workflow depicting the time at which neurons were used following tau oligomer exposure. **B-G** Representative confocal images of immunostaining for presynaptic Synapsin (red) and postsynaptic PSD-95 (green) in human neurons 2 days (**B**), 7 days (**D**), or 14 days (**F**) after exposure to tau monomers or tau oligomers for 30 min. The colocalization of Synapsin and PSD-95 puncta marked synaptic connections on neuronal processes (white arrowheads). Scale bars: 5 μm. The density of individual Synapsin puncta and PSD-95 puncta along neuronal processes (# of puncta/10 µm length) were analyzed to assess presynaptic and postsynaptic compartments separately. The density of colocalized Synapsin and PSD-95 puncta was also measured to assess synapse density (# of puncta/10 µm length). Analyses of puncta densities were performed on neurons that were untreated or treated with tau monomers or tau oligomers for 30 min and then fixed for immunolabeling (**C**) 2, (**E**) 7, and (**G**) 14 days after tau exposure (n = 12 images/group; ** p < 0.01, *** p < 0.001, one-way ANOVA, Bonferroni post hoc analyses) **H** Representative traces from patch clamp recordings of EPSCs in human neurons evoked by a 15 μA extracellular local field stimulation. Each trace represents the mean EPSC of two evoked responses. Human neurons were either exposed to vehicle control (black) or tau oligomers (purple) for 30 min followed by recordings of evoked EPSCs 14 days later. Scale bars: 50 pA, 10 ms. **I** Quantification of the mean amplitude of EPSCs that were evoked in human neurons (n = 24-27 cells/group; * p < 0.05, Unpaired Student’s *t*-test). Values are presented as means ± SEM.

Since the tau oligomer-induced loss of synaptic structures could alter the overall synaptic activity in neurons, we next performed whole-cell voltage clamp recordings on neurons 14 days after acute tau oligomer exposure. To monitor the integrated strength of active synaptic inputs on the neurons, we recorded excitatory postsynaptic currents (EPSCs) elicited by extracellular local field stimulation. Consistent with the loss of functional synaptic inputs, the amplitude of evoked EPSCs was significantly reduced in tau oligomer-treated neurons compared to vehicle-treated controls (Fig. 4H, I). Thus, tau oligomer-induced synaptotoxicity in human neurons involves stages of progressive deterioration over an extended time with the loss of PSD-95-labeled postsynaptic compartments occurring first before the eventual loss of Synapsin-labeled presynaptic terminals which coincides with functional synapse decline.

### Tau oligomers induce a persistent deficit in postsynaptic AMPARs and synaptic plasticity

While the brief tau oligomer exposure caused a progressive loss of synapses, we also wondered how the synapses that remained on the neurons were affected over time. Since tau toxicity is associated with the dysregulation of AMPARs [4], we explored the effect of acute tau oligomer exposure on the levels of postsynaptic AMPARs over time. An antibody to label AMPARs containing the GluA2 and/or GluA3 subunits (GluA2/3) was used to analyze total AMPAR levels, including both surface and intracellular receptors, together with immunolabeling of PSD-95 as a postsynaptic marker. Levels of GluA2/3-containing AMPARs that colocalized with PSD-95 puncta were significantly reduced in neurons at both 7 and 14 days after the acute 30-min tau oligomer exposure compared to vehicle-treated control neurons (Fig. 5A, B, D, E). However, immunostaining of the GluN1 subunit of NMDARs was not altered at synapses of tau oligomer-exposed neurons at either time point (Suppl. Fig. S5A-D), suggesting that total NMDAR levels were maintained. Although the density of PSD-95 puncta, marking the number of postsynaptic sites, was reduced in neurons at 7 days after tau oligomer exposure (Fig. 4D, E), the intensity of PSD-95 immunolabeling at the remaining synapses was not significantly different from controls (Fig. 5C). This indicates that, despite tau oligomer-induced loss of postsynaptic sites at 7 days, the PSD-95 levels in remaining synapses were intact. Yet, by 14 days after tau oligomer exposure, the intensity of PSD-95 immunolabeling was significantly reduced (Fig. 5F), signifying a progressive overall weakening of the remaining postsynaptic compartments over time. These data suggest that the acute tau oligomer exposure causes a long-term downregulation of AMPARs, but not NMDARs, and a structural decline at the postsynaptic sites that remained. Since tau oligomers acutely blocked synaptic plasticity (Fig. 1A), we tested whether synaptic plasticity was still disrupted 14 days after the 30-min tau oligomer exposure in the remaining synapses. Surprisingly, the insertion of surface postsynaptic GluA1-containing AMPARs after cLTP induction was blocked in neurons exposed to tau oligomers 14 days prior, whereas vehicle-treated control neurons maintained cLTP-induced AMPAR delivery (Fig. 5G, H). These results support that tau oligomers not only cause synapse loss, but they trigger persistent postsynaptic dysfunction that lasts for weeks after exposure in remaining synapses.

**Figure 5.**
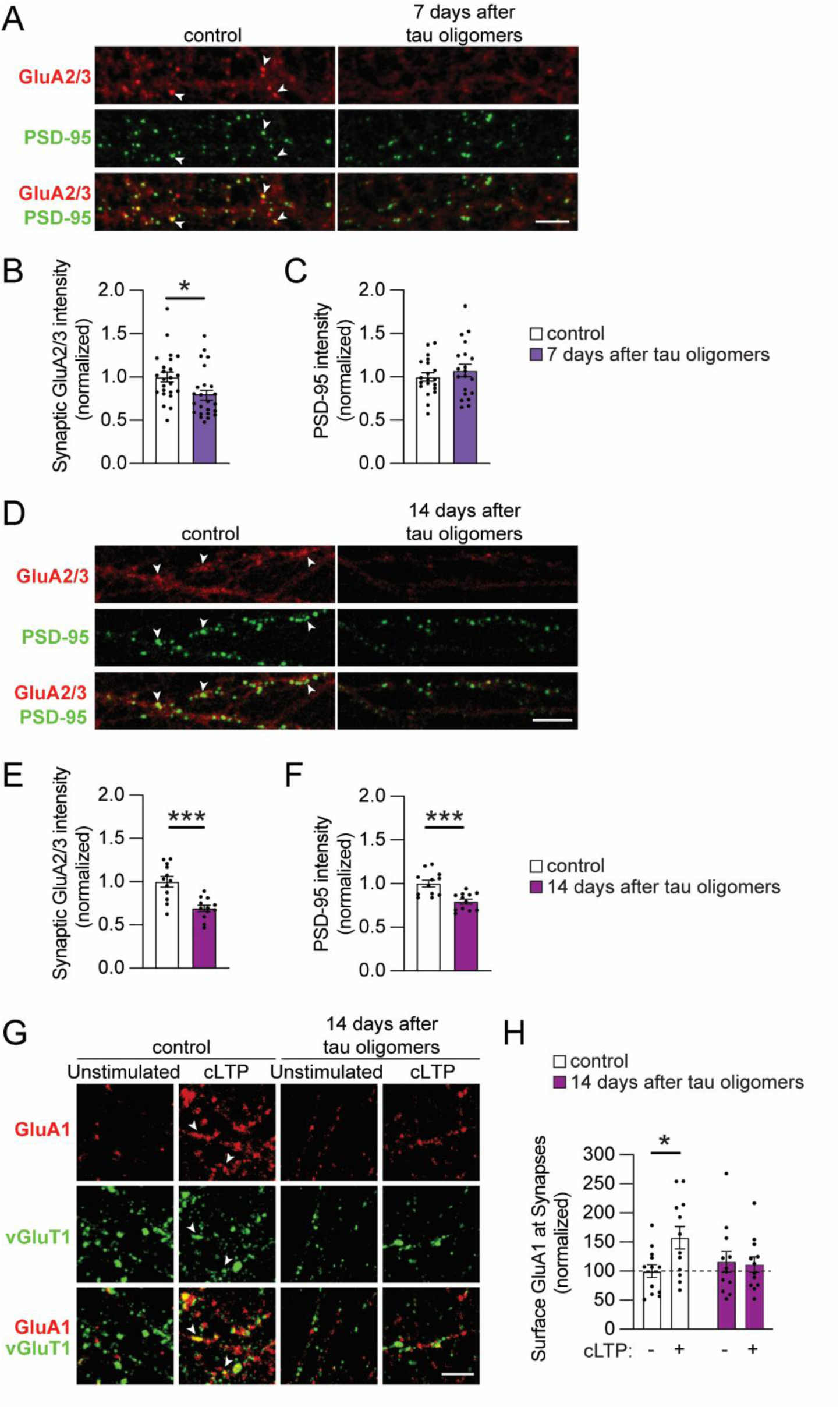
Tau oligomer exposure induced lasting deficits in postsynaptic AMPARs and LTP expression in human neurons. A-F. Representative confocal images of immunostaining for the total levels of GluA2/3 subunit-containing AMPARs (red), including both surface and intracellular AMPAR pools, and PSD-95 (green) in human neurons at (**A**) 7 or (**D**) 14 days after the acute exposure to tau oligomers and vehicle-treated control neurons. The mean intensity of GluA2/3 immunofluorescence that colocalized with PSD-95 puncta was quantified (arrowheads) at (**B**) 7 or (**E**) 14 days post-tau oligomer exposure and normalized to vehicle-treated control neurons. The mean intensity of PSD-95 puncta was also quantified at (**C**) 7 or (**F**) 14 days post-tau oligomer exposure normalized to vehicle-treated controls (n = 12-24 images/group; * p < 0.05, *** p < 0.05, Unpaired Student’s *t*-test). **G** Representative confocal images of surface GluA1-containing AMPARs labeled with a GluA1 antibody (red) that co-localized with vGluT1 (green), a synaptic marker, in human neurons (arrowheads). Neurons were treated with vehicle or tau oligomers for 30 min, and then 14 days later, the neurons were either unstimulated or stimulated by cLTP induction. Scale bar: 5 μm. **H** Graph of the immunolabeling intensity of surface GluA1 quantified at synapses 14 days after tau oligomer treatment. The surface GluA1 intensities across groups were normalized to unstimulated control neurons (n = 12 neurons/group; * p < 0.05 Unpaired Student’s *t*-test).

### Tau oligomers disrupt the localization and release of synaptic vesicles from presynaptic terminals

Our findings demonstrate that tau oligomers cause a long-term dysregulation of postsynaptic function, which could occur simultaneously with presynaptic changes. Notably, the expression of tau carrying mutations that cause familial frontotemporal dementia inhibits presynaptic function by hindering synaptic vesicle mobility [103, 104]. To investigate the lasting impact of the 30-min tau oligomer exposure on presynaptic terminals in human neurons, electron microscopy was performed for ultrastructural analyses of the presynaptic terminals that remained 7 and 14 days after the exposure (Fig. 6A). Intriguingly, at 7 and 14 days after acute tau oligomer exposures, neurons had significantly increased numbers of synaptic vesicles within each presynaptic terminal compared to vehicle-treated control neurons (Fig. 6B). However, unlike control neurons that had clusters of vesicles assembled around the active zones in each terminal, most terminals in tau oligomer-treated neurons did not have any clustered vesicles (Fig. 6C). Vesicle clustering at the active zone is critical for both the structure and function of synapses [105]. Thus, the accumulation of highly disordered synaptic vesicles without proper clustering could have a multifaceted impact on presynaptic function.

**Figure 6.**
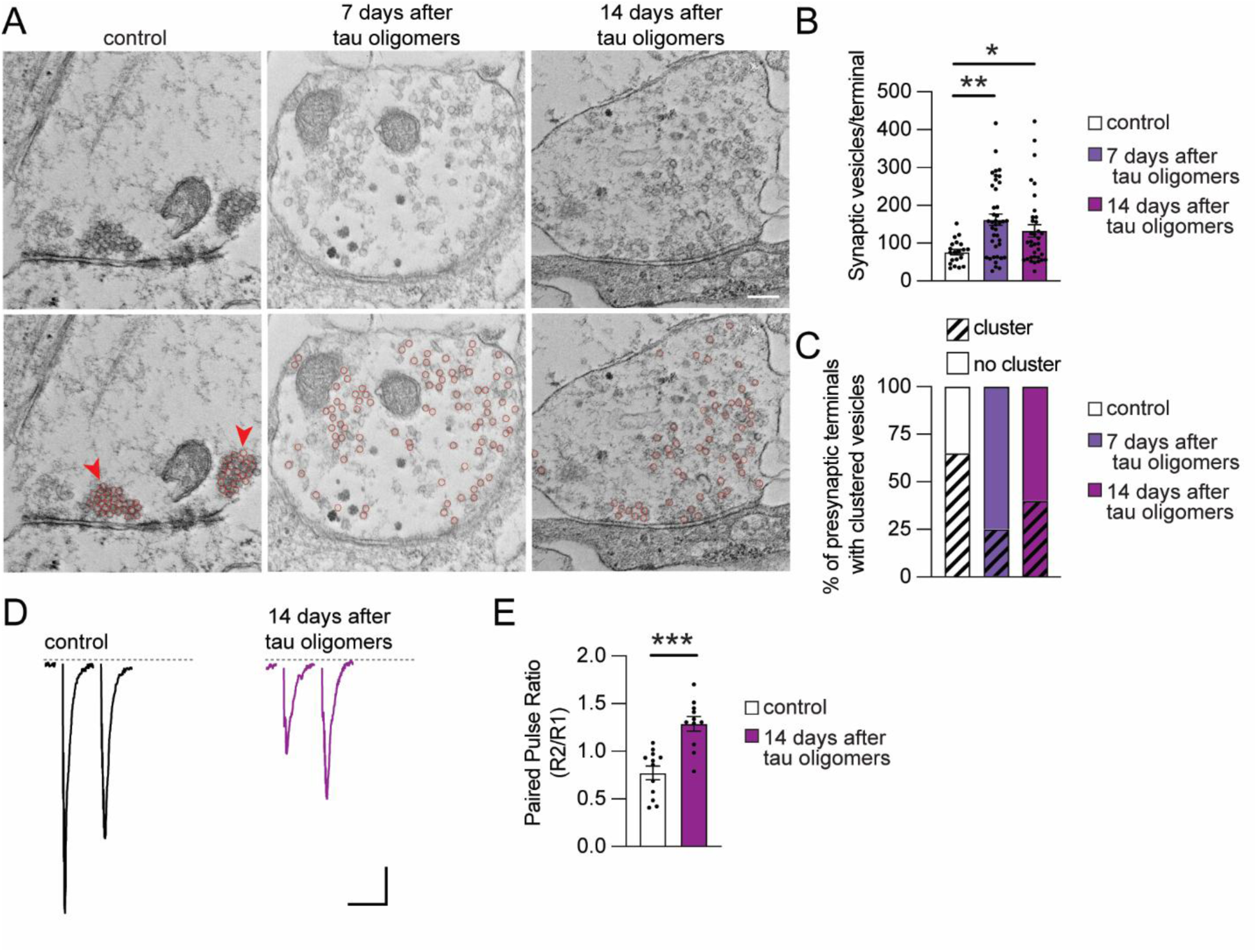
Long-term effect of acute tau oligomer exposure on presynaptic vesicle organization and vesicle release probability. **A** Representative electron microscopy images of human neuron synapses 7 or 14 days after treatment with either tau oligomers for 30 min or the vehicle control (top). The respective identical images below depict the synaptic vesicles highlighted by red circles. The synaptic vesicles in the vehicle-treated control presynaptic terminal are clustered close to the active zone (red arrows). Scale bar: 200 nm. Magnification: 50,000 x. **B** Graph of the number of synaptic vesicles quantified in each presynaptic terminal of the human neurons with and without tau oligomer exposure (n = 20-40 synapses/group; one-way ANOVA, Bonferroni post hoc analyses). **C** Graph of the percentage of presynaptic terminals at human neuron synapses calculated that contained vesicle clusters (number of terminals with clustered vesicles/total synapses; control: 13/20; 7 days: 10/40; 14 days: 14/35). **D** Representative traces from patch clamp recordings of EPSCs in human neurons elicited by two consecutive extracellular local field stimuli (15 μA) with a 20 ms interstimulus interval (ISI). Recordings were performed 14 days after human neurons were either exposed to vehicle control (black) or tau oligomers (purple) for 30 min. Scale bars: 50 pA, 20 ms. **E** The paired pulse ratio (PPR) was calculated at the 20 ms ISI by dividing the amplitude of the second evoked EPSC (R2) by the amplitude of the first evoked EPSC (R1) (n = 11-12 cells/group; Unpaired Student’s *t*-test).

To determine whether synaptic vesicle release was altered by tau oligomer exposure, whole-cell patch-clamp recordings were performed to measure the paired-pulse ratio (PPR) from consecutive evoked EPSCs, which indicates the synaptic vesicle release probability (Fig. 6D). The PPR is calculated for each neuron by dividing the amplitude of the second EPSC response (R2) by the amplitude of the first EPSC response (R1). The PPR measured from neurons 14 days after the 30-min tau oligomer exposure was significantly increased compared to control neurons (Fig. 6E), revealing that tau oligomers cause a long-term decrease in synaptic vesicle release probability at the remaining synapses. Together, these data link the disordered localization of synaptic vesicles at presynaptic terminals with a lower vesicle release probability at the dysfunctional synapses that remain up to 14 days after tau oligomer exposure.

## Discussion

Our study elucidates how a brief tau oligomer exposure initiates a progressive cascade of events that lead to either bipartite synapse dysfunction or synapse loss (Fig. 7). Interestingly, the emergence of tau-mediated presynaptic and postsynaptic pathophysiological changes occurred in parallel as synapses deteriorated. Tau oligomers caused a long-term disorganization of synaptic vesicles at presynaptic terminals on human neurons which coincided with the downregulation of GluA2/3-containing AMPARs at postsynaptic sites. Moreover, tau oligomers caused both an immediate and a long-standing inhibition of LTP expression at synapses, indicating a persistent impairment of plasticity mechanisms. Temporal mapping of the postsynaptic proteome revealed distinct time-dependent alterations in postsynaptic proteins after tau oligomer exposure, indicating the dynamic nature of the processes that underlie synapse deterioration.

**Figure 7.**
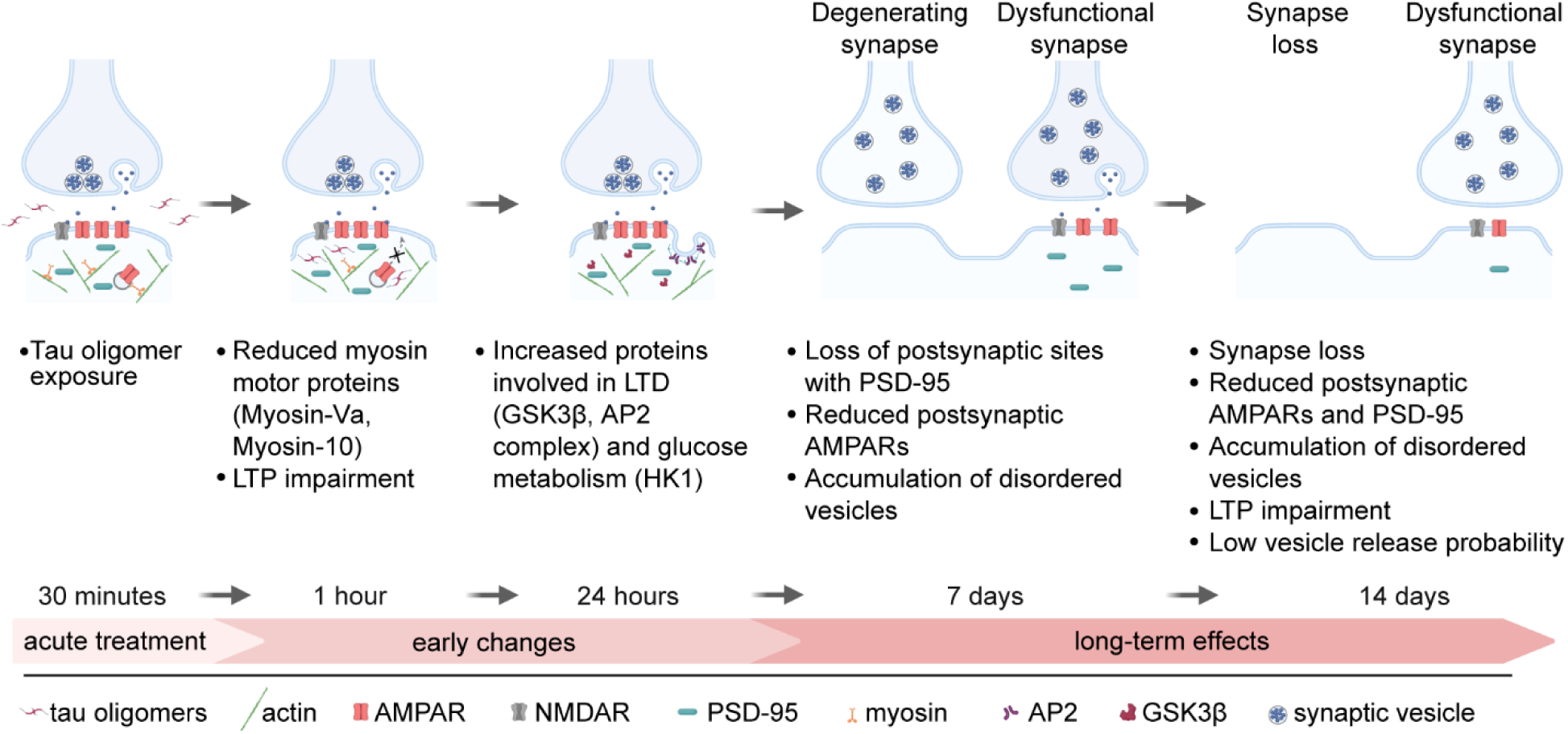
Effect of tau oligomers on synapse fate. Illustration depicting the series of events during progressive synapse decline that follows a brief exposure of human neurons to tau oligomers. Within 1 hour after exposure, reduced postsynaptic localization of myosin motor proteins coincides with impaired postsynaptic AMPAR insertion for LTP expression. One day after exposure, disease-related proteins involved in LTD expression and glucose metabolism were upregulated at postsynaptic sites. At 7 days after tau oligomer exposure, presynaptic terminals contain disordered synaptic vesicles and some postsynaptic compartments containing PSD-95 are lost while the postsynaptic sites that remain contain less AMPARs. At 14 days after tau oligomer exposure, some bipartite synapses comprised of both presynaptic and postsynaptic compartments are lost, while other synapses persist with aberrant presynaptic vesicle release and impaired postsynaptic AMPAR regulation.

Postsynaptic proteome mapping revealed reduced postsynaptic Myosin-Va levels after tau oligomer exposure that was associated with the immediate inhibition of LTP expression at synapses. Myosin-Va is a Ca^2+^-dependent actin-based motor protein that transports intracellular cargoes [106]. Importantly, Myosin-Va is required for LTP expression because it mediates the activity-dependent transport of AMPARs into postsynaptic spines in mouse hippocampal neurons [64]. This raises the possibility that the rapid downregulation of postsynaptic Myosin-Va by tau oligomers inhibits AMPAR trafficking for LTP expression. Myosin-Va also regulates the transport of NMDARs [65], but we did not detect an overall change in postsynaptic NMDAR levels after tau oligomer treatment. Myosin-10, another postsynaptic motor protein that was reduced by tau oligomers, promotes the formation of filopodia and synapses on dendrites through actin cytoskeletal-based mechanisms [67, 107]. Intriguingly, Myosin-Va and Myosin-10 were identified as tau-interacting proteins in tau interactome analyses [59, 108]. Whether the internalization and transport of tau oligomers at synapses is related to the interaction between tau and the actin-based motor proteins or the downregulation of Myosin-Va and Myosin-10 at postsynaptic sites remains to be investigated. A fundamentally different effect was detected in the postsynaptic proteome 24 hours after tau oligomer exposure, namely the upregulation of disease-related postsynaptic proteins. These findings support a time-dependent emergence of distinct mechanisms that drive synapse pathophysiology. Notably, the upregulation of postsynaptic GSK3β 24 hours after tau oligomer exposure could cause both impaired LTP and enhanced LTD at synapses [86, 91]. Moreover, GSK3β phosphorylates 29 residues on the tau protein [83, 87, 109, 110], which is associated with the weakening of synapse function and AD-related pathogenesis [84–87]. Consistent with our findings, tau oligomers increase GSK3β levels in mouse hippocampus underlying tau hyperphosphorylation and memory impairment [111]. Mitochondrial dysfunction has also been linked to synapse loss triggered by tau oligomers and contributes to the formation of tau oligomers [13, 112–114]. We detected increased postsynaptic levels of mitochondrial proteins related to glucose metabolism 24 hours after tau oligomer exposure. These mitochondrial proteins could modulate energy consumption in neurons, which is dysregulated in AD [115]. Two of the mitochondrial proteins identified, SLC25A6 and CS, were identified as tau-interacting proteins with reduced binding to disease-related tau mutants linked to mitochondrial dysfunction [59].

One day after the brief exposure, tau oligomers were no longer detectable in the human neurons, suggesting that other mechanisms initially triggered by the tau oligomers drive the longer-term effects on synapses. Exogenous tau oligomers can be internalized by neurons and induce the seeding of pathological endogenous tau proteins [19, 24, 116, 117]. It is possible that the delayed MAP2-associated dendritic cytoskeletal deterioration and impaired synaptic transmission we observed occurred due to the seeding of pathogenic endogenous tau or due to the accumulated damage at synapses triggered by the initial internalization of tau oligomers. In addition to synaptotoxicity, the uptake of oligomeric tau has also been linked to the formation of stress granules [25, 118–120], nuclear abnormalities [121], axonal transport dysregulation [116], axon initial segment degeneration [122], and mitochondrial dysfunction [13]. Similar to previous reports [36], the effect of tau oligomers on synapse deterioration did not coincide with neuron loss in our study, but depending on the experimental conditions, tau oligomers may drive more severe neurodegenerative phenotypes [27].

How are synapses lost during tau-mediated pathogenesis? Our findings suggest that synapses undergo a progressive deterioration following tau oligomer exposure that involves changes in both presynaptic and postsynaptic compartments. Interestingly, the human neurons lost postsynaptic sites, marked by PSD-95, before they lost Synapsin-containing presynaptic terminals, which suggests that the bipartite synaptic connections are not necessarily lost simultaneously. Moreover, this progressive weakening of overall synapse function is consistent with the notion of intrinsic synaptic signaling pathways causing synapse elimination after LTD induction [123, 124]. Indeed, a brief tau oligomer exposure promotes LTD expression in hippocampus [34]. On the other hand, synapses can be engulfed by microglia or astrocytes in tauopathy mouse models in a maladaptive process that is similar to synaptic pruning [125–127]. A recent study on postmortem human brain tissue reported an increase in tau oligomer-containing synapses that colocalized with astrocytes and microglia in AD brains [18], raising the possibility that glia also play a role in tau oligomer-mediated synapse loss. The human neurons in our study were co-cultured with astrocytes, but the cultures lacked microglia. The extent to which synapse elimination triggered by pathogenic tau is driven by intrinsic synaptic weakening compared to glial-mediated synapse engulfment under different pathological conditions remains unclear. Yet our findings show that tau oligomer-mediated synapse dysfunction and synapse loss are both triggered by a chain of events over two weeks involving both presynaptic and postsynaptic dysregulation.

## Conclusion

Overall, our results reveal the timeline of pathophysiological events triggered by tau oligomers at synapses of human neurons leading to the elimination of some synapses and the prolonged presynaptic and postsynaptic dysfunction at remaining synapses. Immediately after tau oligomer exposure, the downregulation of postsynaptic myosin motors overlapped with impaired postsynaptic AMPAR insertion after LTP induction. The subsequent increase in levels of postsynaptic GSK3β and other disease-related proteins was followed by a loss of postsynaptic and presynaptic sites 7 days and 14 days after tau oligomer exposure, respectively. Synapses that persisted after tau oligomer exposure had reduced postsynaptic PSD-95 and AMPAR levels, impaired LTP expression, and presynaptic terminals with accumulated un-clustered synaptic vesicles and lower vesicle release probability.

## Acknowledgements

We thank the Buck Institute Proteomics and Mass Spectrometry Core, Stella Breslin and Harris Ingle from the Buck Institute Morphology and Imaging Core, and Lee Cohen-Gould from the Weill Cornell Medicine Microscopy and Image Analysis Core for support services. We thank Mary Redwine for administrative support. We are grateful to Dr. Li Gan (Weill Cornell Medicine) for the human iPSCs and lentivirus plasmids and Dr. Sue-Ann Mok (University of Alberta) for the human tau 2N4R plasmid.

## Author contributions

K.A.P-N. and T.E.T. conceived the project and designed experiments. K.A.P-N., C.D.K., G.K., D.L., A.V., and T.E.T performed experiments. Y.Y.N., I.W., J.H.C, M.S. and G.N. provided technical support. K.A.P-N. and T.E.T. wrote the manuscript. All authors reviewed the manuscript.

## Abbreviations

iPSC: Induced pluripotent stem cell
AD: Alzheimer’s disease
APEX2-PSD-95: APEX2 fused with PSD-95
NGN2: Neurogenin 2
AMPAR: a-amino-3-hydrocy-5-methyl-4-isoxazolepropionin acid receptor
NMDAR: N-methyl-D-aspartate receptor
LTP: Long-term potentiation
LTD: Long-term depression
PPR: Paired-pulse ratio
SynGO: Synaptic gene ontologies
FITC: Fluorescein-5-isothiocyanate
PFA: Paraformaldehyde
GSEA: Gene set enrichment analysis
cLTP: Chemical long-term potentiation
EPSC: Excitatory postsynaptic currents
FBS: Fetal bovine serum

