## Supplementary figures for "Tau oligomers modulate synapse fate by eliciting progressive bipartite synapse dysregulation and synapse loss"

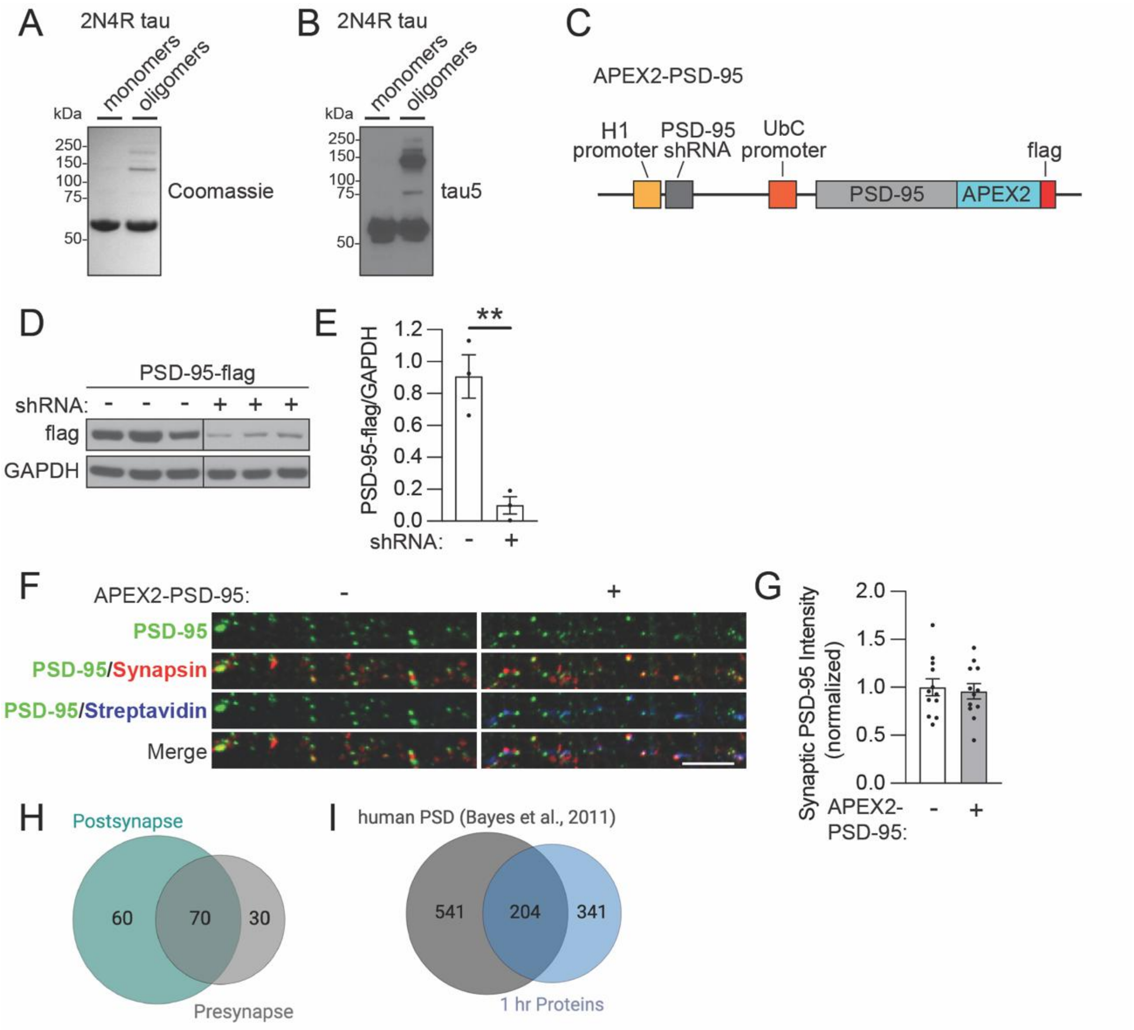


**Supplemental Figure 1: Characterization of recombinant human tau oligomers and the APEX2-PSD-95 strategy of postsynaptic mapping.**

(**A**) Non-reducing SDS-PAGE gel of recombinant human 2N4R tau monomers and tau oligomers preparations stained with Coomassie dye.

(**B**) Immunoblot with a tau5 antibody on recombinant human 2N4R tau monomers and tau oligomers preparations.

(**C**) Design of the APEX2-PSD-95 construct containing a ubiquitin C promoter (UbC) to drive APEX2-PSD-95-flag expression and an H1 promoter to drive the expression of a short hairpin RNA (shRNA) to knockdown endogenous human PSD-95.

(**D**) Immunoblots of lysates from HEK293 cells expressing human flag-tagged PSD-95 with or without PSD-95 shRNA.

(**E**) Quantification of PSD-95-flag immunolabeling on the western blot from HEK293 cell lysates with or without PSD-95 shRNA expression normalized to GAPDH levels (n = 3 wells/treatment; ** $p < 0.01$, Unpaired Student's *t*-test)

(**F**) Representative images of PSD-95 (green) colocalized with Synapsin (red) in untransfected control neurons and neurons expressing APEX2-PSD-95. Streptavidin (blue) staining shows neurons that have APEX2-PSD-95-mediated biotinylation. Scale bar: 5 µm

(**G**) Quantification of PSD-95 intensity levels co-localized with Synapsin in uninfected control neurons and neurons expressing APEX2-PSD-95. Values were normalized to the mean PSD-95 intensity in the uninfected control neurons (n = 12 images/group, not significant, Unpaired Student's *t*-test).

(**H**) Venn diagram of biotinylated proteins categorized in postsynapse and presynapse SynGo cellular component analysis showed an overlap between 70 proteins in both presynaptic and postsynaptic compartments.

(**I**) Venn diagram revealed the overlap between the 545 biotinylated proteins detected in human neurons with APEX2-PSD-95-mediated proximity labeling and 745 proteins identified in the human postsynaptic density by mass spectrometry performed by Bayes et al., 2011.

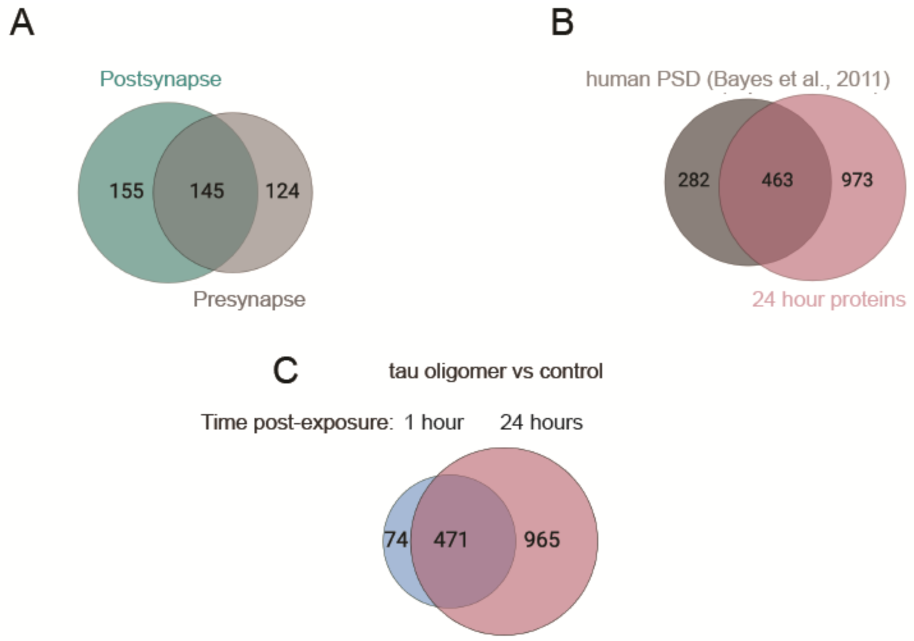


**Supplemental Figure 2: Functional relationship and characterization of biotinylated proteins identified by mass spectrometry 24 hours after exposure of human neurons to tau oligomers.**
(**A**) Venn diagram of annotated genes from SynGo cellular component analysis showing overlap between 300 postsynaptic genes and 269 presynaptic proteins.
(**B**) Venn diagram showing overlap between 1436 proteins identified by mass spectrometry 24 hours after exposure of human neurons to tau oligomers and 745 proteins from human postsynaptic density identified by mass spectrometry performed by Bayes et al., 2011.
(**C**) Venn diagram showing overlap between 545 and 1436 proteins identified by mass spectrometry 1 hour and 24 hours after exposure of human neurons to tau oligomers.

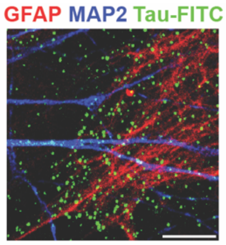


**Supplemental Figure 3: Internalization of tau oligomers by astrocytes in the human neuron cultures.** Confocal image of human neurons treated with oligomerized tau-FITC for 30 min show colocalization of tau-FITC (green) with the astrocytic marker, GFAP (red). Scale bar: 10 μm.

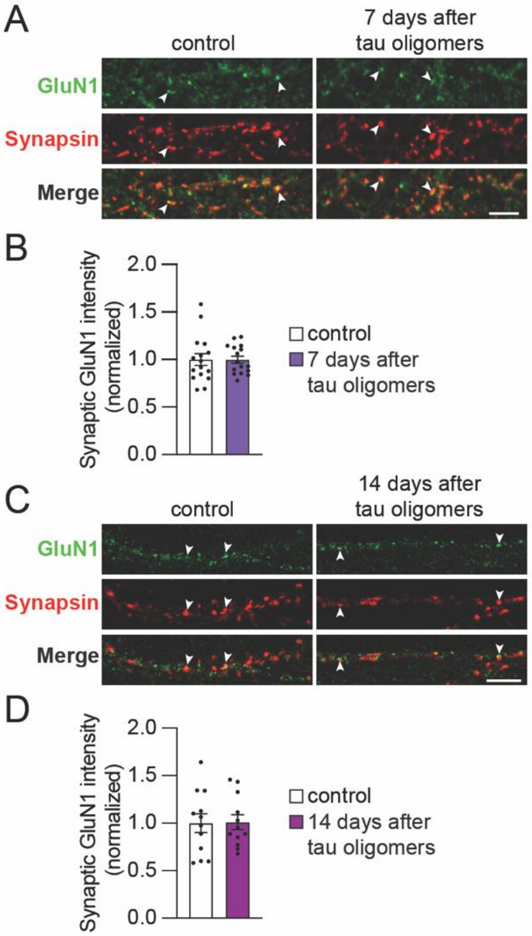


**Supplemental Figure 4: Acute treatment with tau oligomers does not alter NMDAR levels at synapses in human neurons.**

(**A, B**) Representative confocal images (**A**) and quantification (**B**) of GluN1 (green) colocalized with the presynaptic marker Synapsin (red, arrowheads) in human neurons 7 days after exposure to tau oligomers for 30 min (n = 16 images/group; no significant difference, Unpaired Student's *t*-test). Scale bar: 5 µm.

(**C, D**) Representative confocal images (**C**) and quantification (**D**) of GluN1 (green) colocalized with the presynaptic marker Synapsin (red) in human neurons 14 days after exposure to tau oligomers for 30 min (n = 12 images/group; no significant difference, Unpaired Student's *t*-test). Scale bar: 5 µm.
